# Genome wide association study pinpoints key agronomic QTLs in African rice *Oryza glaberrima*

**DOI:** 10.1101/2020.01.07.897298

**Authors:** Philippe Cubry, Hélène Pidon, Kim Nhung Ta, Christine Tranchant-Dubreuil, Anne-Céline Thuillet, Maria Holzinger, Hélène Adam, Honoré Kam, Harold Chrestin, Alain Ghesquière, Olivier François, François Sabot, Yves Vigouroux, Laurence Albar, Stefan Jouannic

## Abstract

**Background:** African rice, *Oryza glaberrima*, is an invaluable resource for rice cultivation and for the improvement of biotic and abiotic resistance properties. Since its domestication in the inner Niger delta ca. 2500 years BP, African rice has colonized a variety of ecologically and climatically diverse regions. However, little is known about the genetic basis of quantitative traits and adaptive variation of agricultural interest for this species.

**Results:** Using a reference set of 163 fully re-sequenced accessions, we report the results of a Genome Wide Association Study carried out for African rice. We investigated a diverse panel of traits, including flowering date, panicle architecture and resistance to *Rice yellow mottle virus*. For this, we devised a pipeline using complementary statistical association methods. First, using flowering time as a target trait, we demonstrated that we could successfully retrieve known genes from the rice flowering pathway, and identified new genomic regions that would deserve more study. Then we applied our pipeline to panicle- and resistance-related traits, highlighting some interesting QTLs and candidate genes (including *Rymv1* for resistance and *SP1*, *Ghd7.1*, *APO1* and *OsMADS1* for panicle architecture). Lastly, using a high-resolution climate database, we performed an association analysis based on climatic variables, searching for genomic regions that might be involved in adaptation to climatic variations.

**Conclusion:** Our results collectively provide insights into the extent to which adaptive variation is governed by sequence diversity within the *O. glaberrima* genome, paving the way for in-depth studies of the genetic basis of traits of interest that might be useful to the rice breeding community.

## BACKGROUND

African rice, *Oryza glaberrima* Steud., was domesticated independently of Asian rice *Oryza sativa* L. (Wang et al. 2014; Meyer et al. 2016; Cubry et al. 2018; Choi et al. 2019). Its domestication took place in the inner delta of the Niger river (Cubry et al. 2018), from a wild relative species, *Oryza barthii* A. Chev.. Its origin from this wild Sahelian species certainly explains its strong tolerance or resistance to biotic and abiotic stresses (Sarla and Swamy 2005). In the context of increasing temperatures and a more variable climate, strong tolerance to such stresses is an important objective for rice agriculture worldwide. However, knowledge of the genetic basis of phenotypic variation in African rice remains very limited. With the exception of salinity tolerance (Meyer et al. 2016), few association studies have been performed for traits of agricultural interest in this species. Genome wide association studies (GWAS) have successfully identified genes of functional importance associated with flowering time in Asian rice (Zhao et al. 2011; Huang et al. 2012; Yano et al. 2016). For Asian rice, the genetic determination of this trait is well understood (Lee and An 2015), whereas we have no information about the variation of this trait for African rice. Another trait of broad interest for rice farmers and breeder communities is the architecture of the panicle. This trait is one of the main components of yield potential, because the number of seeds per panicle is directly related to the branching complexity of the inflorescence (Xing and Zhang 2010). With increasing global movement of plant material and climate change, biotic threats to rice agriculture continue to evolve and the search for new sources of resistance to pathogens is therefore a challenging research field. *Rice yellow mottle virus* (RYMV) is responsible for one of the most damaging diseases of rice in Africa (Kouassi et al. 2005; Issaka et al. 2012; Kam et al. 2013). Resistance genes against RYMV are mostly found in *O. glaberrima*, and this species may be an interesting source of quantitative trait loci (QTLs) for global rice breeding strategies (Thiémélé et al. 2010).

To better assess the functional variation present in African rice, we developed a genome-wide association panel and corresponding phenotypic datasets for flowering time, inflorescence architecture, and resistance to RYMV. Using several complementary statistical models for genetic association, we identified key QTLs for flowering time variation, panicle architecture, quantitative resistance to RYMV and climatic variation.

## RESULTS

A total of 892,539 SNPs from 163 different *O. glaberrima* accessions was used in the GWAS. We considered that population structure was best described by four genetic groups, and we used either one or both of the groupings and the kinship matrix as cofactors to correct for confounding effects in our analyses. The phenotypic data were obtained from infield experiments or from available public databases (data in Additional file 1: Table S1). Using ANOVA as the benchmark, all methods allowed an efficient correction for false positives (QQ-plots, see Additional files 2, 3, 4, 5).

### Flowering time

A genome-wide association study of flowering time based on data from the early planting dates allowed us to identify 1,450 SNPs statistically associated with this trait (5% FDR threshold, Table 1; Additional file 2; Additional file 6: Table S2). Most of these SNPs were at a distance of less than 25kb from each other and were clumped into 80 genomic regions containing a total of 733 annotated genes (Table 1; Additional file 7: Table S3; Additional file 8: Table S4).

**Table 1.** Numbers of significant SNPs, regions and genes found to be associated with the different traits. Significant SNPs were detected with the different models using the Fisher combination method and based on a FDR 5% threshold. The "All methods" column indicates the number of SNPs detected with at least one method. Fifty kb windows around these SNPs defined independent genomic regions associated with each trait.

| Trait | Significant SNPs |  |  |  |  | Regions | Genes |
| --- | --- | --- | --- | --- | --- | --- | --- |
|  | EMMA | GAPIT | LFMM | CATE | All methods |  |  |
| **BioPC1** | 0 | 0 | 0 | 0 | 0 | 0 | 0 |
| **BioPC2** | 47 | 0 | 38 | 42 | 52 | 17 | 128 |
| **TmaxPC1** | 0 | 0 | 0 | 0 | 0 | 0 | 0 |
| **TmaxPC2** | 2,308 | 78 | 1,559 | 1,656 | 2,378 | 228 | 1,946 |
| **Early Sowing** | 850 | 0 | 1,343 | 1,044 | 1,450 | 80 | 732 |
| **Late Sowing** | 251 | 0 | 2,042 | 2,535 | 2,649 | 134 | 1,223 |
| **RYMV** | 2,171 | 387 | 197 | 156 | 2,264 | 257 | 2,248 |
| **SpN** | 0 | 0 | 0 | 0 | 0 | 0 | 0 |
| **PBN** | 0 | 0 | 0 | 257 | 257 | 25 | 243 |
| **SBN** | 1 | 0 | 0 | 0 | 1 | 1 | 10 |
| **RL** | 243 | 0 | 0 | 0 | 243 | 38 | 361 |
| **PBL** | 0 | 0 | 56 | 54 | 57 | 12 | 91 |
| **PBintL** | 1 | 0 | 0 | 1 | 1 | 1 | 9 |
| **SBintL** | 123 | 0 | 281 | 454 | 455 | 30 | 266 |

Corresponding analyses performed using data from the later planting date revealed associations with a slightly larger number of significant SNPs: 2,649 significant SNPs corresponding to 134 regions and 1,223 annotated genes (Table 1; Additional file 2; Additional file 6: Table S2; Additional file 7: Table S3; Additional file 8: Table S4). Two hundred and sixty-two genes representing 25 genomic regions were found for both planting dates. Several GWAS peaks co-localized with known Asian rice flowering time genes either for early or late planting (Table 2; Fig. 1): *OsGI* and *OsMADS51* on chromosome 1; *Hd6*/*OsMADS14*/*OsPHYC* and *ETR2* on chromosome 3; *Hd3a*/*RFT1* on chromosome 6; and *RCN1* on chromosome 11. When we compared the list of genes found in our study for flowering time with an expert list based on bibliography (Additional file 9: Table S5), we found a five-fold significant enrichment (G test associated *p*-value = 7e10^−4^), showing that our GWAS approaches were effective in retrieving known genes.

**Table 2.**
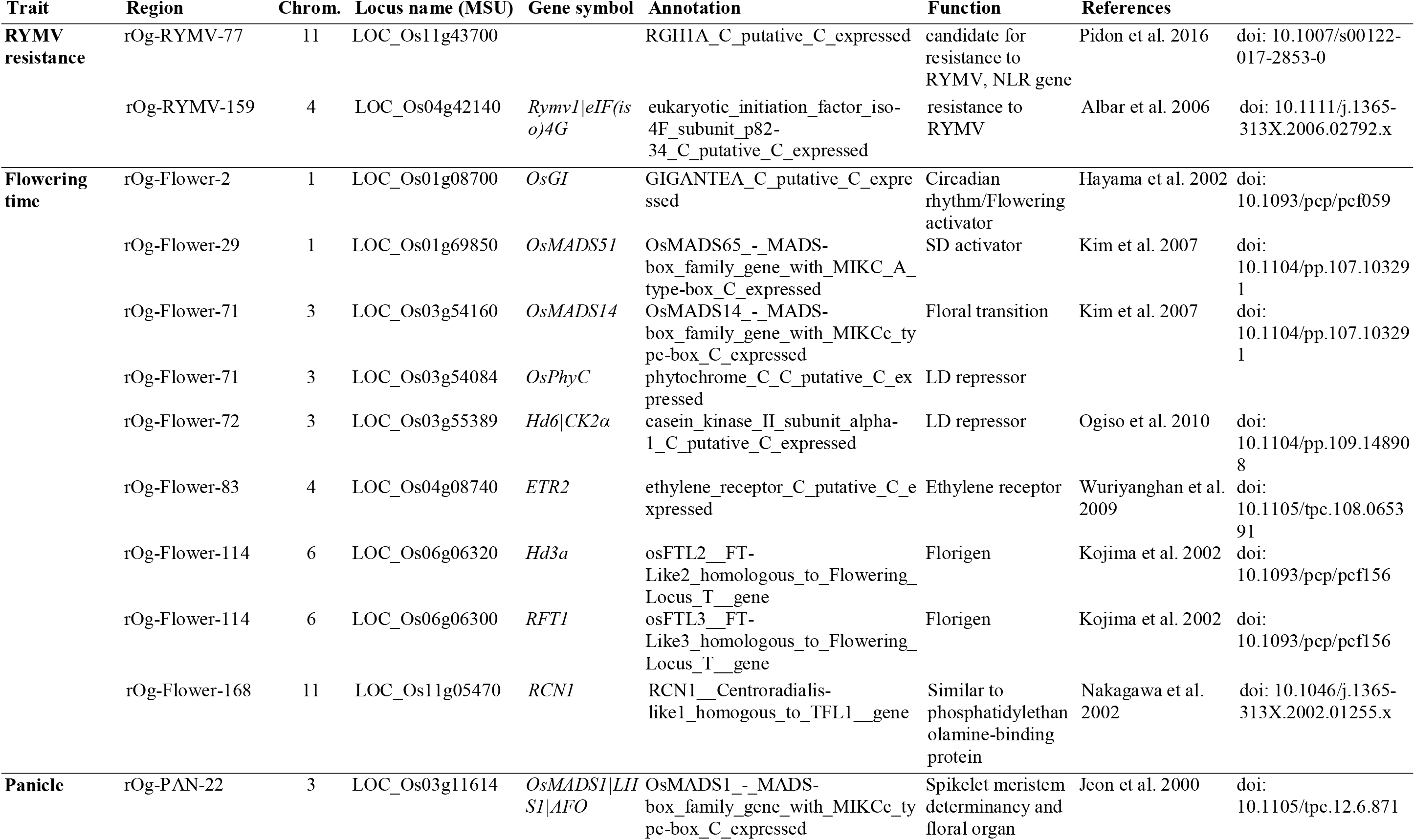
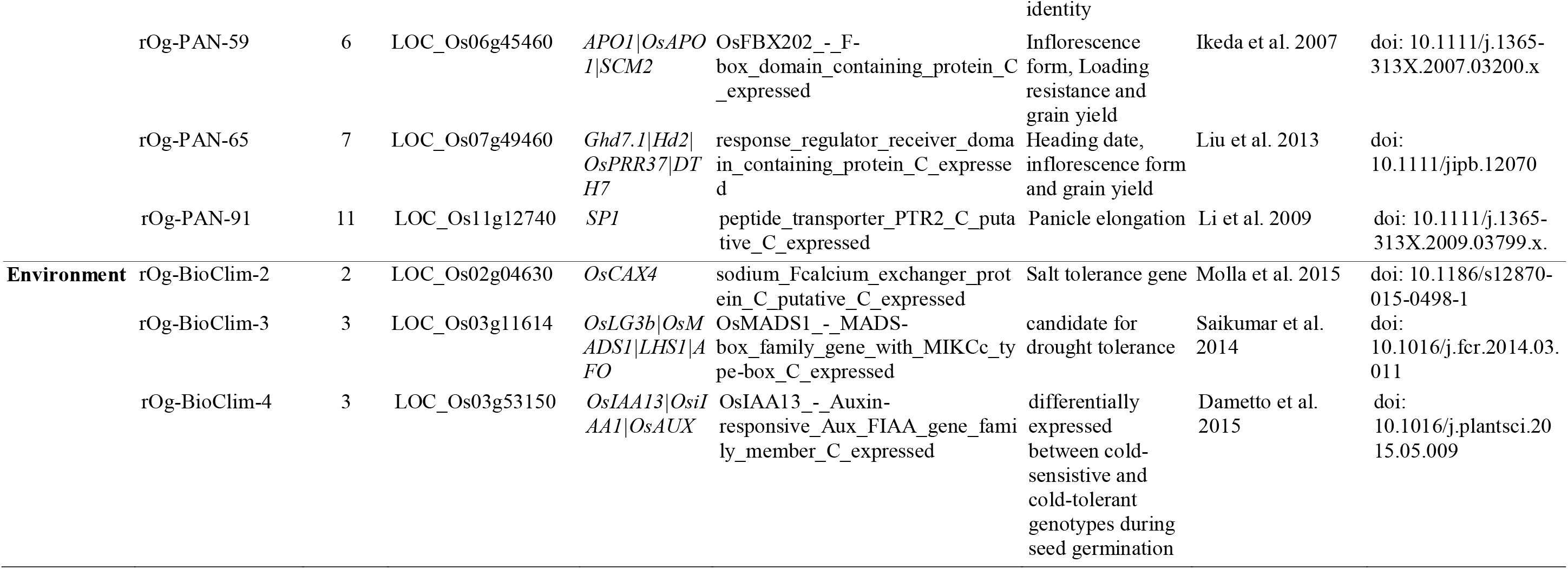
List of the main candidate genes identified in regions associated with RYMV resistance, flowering time, panicle architecture and climate variation.

**Figure 1.**
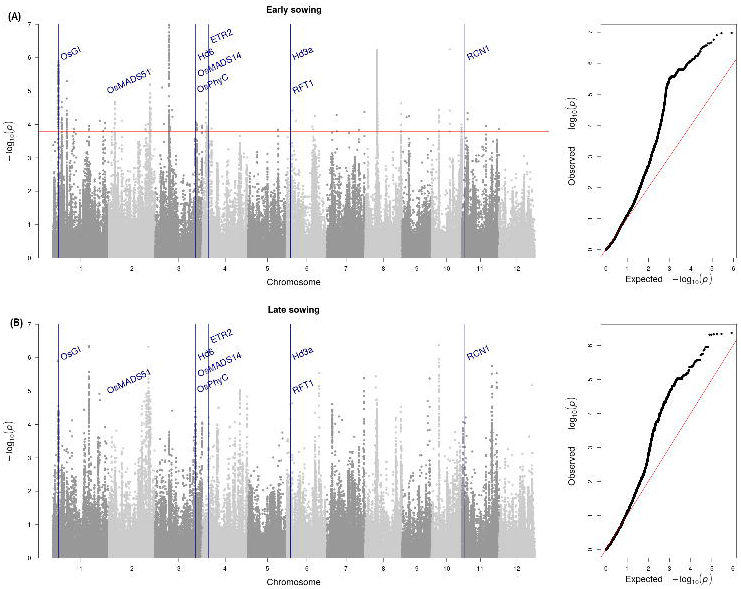
Manhattan plots of LFMM association results for flowering time assessed for early (upper pane) and late (lower pane) sowing. The red line indicates the 5% FDR threshold. Known Asian rice flowering genes found in the vicinity of significant SNPs are indicated.

GO term enrichment analysis performed on the complete list of genes found five significantly (p-value < 0.05) enriched GO terms using classic Fisher and weighted tests, two for cellular components (Golgi apparatus and peroxisome) and three for molecular functions (motor activity, carbohydrate binding and transcription factor activity) (Additional file 10: Table S6). An additional GO term was found for biological processes (Cellular processes) with the weighted test only.

### Panicle morphological traits

Using six morphological traits, all association methods led to comparable numbers of significant associations (Table 1). The CATE model produced the highest number of significant SNPs for four of the six morphological traits. A total of 1,010 significant SNPs was detected for at least one panicle morphological trait, except for spikelet number for which no significant SNP was found (Table 1; Additional file 3; Additional file 6: Table S2). Ninety-seven unique genomic regions associated with one or more morphological traits were defined (Table 1; Additional file 7: Table S3). Nine of the 97 unique regions were associated with more than one trait: one for secondary branch number (SBN) and primary branch average length (PBL) (rOg-PAN-18), three for primary branch number (PBN) and rachis length (RL) (rOg-PAN-25, rOg-PAN-58, rOg-PAN-92), two for PBN and secondary branch internode average length (SBintL) (rOg-PAN-8, rOg-PAN-56), two for RL and SbintL (rOg-PAN-78, rOg-PAN-88) and one for PBN, RL and SBintL (rOg-PAN-90). The 88 remaining regions were each associated with a single trait: 32 for RL, 20 for PBN, 10 for PBL, 1 for primary branch internode average length (PBintL) and 25 for SBintL. Among the 97 associated regions, 858 annotated genes were identified (Table 1; Additional file 8: Table S4). GO term analysis revealed no significant enrichment of gene categories in the regions of significant SNPs (Additional file 10: Table S6).

Some genes already known to be involved in panicle development control co-localized with genomic regions associated with panicle morphological traits, examples including: *ABERRANT PANICLE ORGANIZATION1* (*APO1*) which colocalizes with the region rOg-PAN-59 identify for the SBintL trait; *SHORT PANICLE1* (*SP1*) with the rOg-PAN-91 region (PBL); *OsMADS1*/*LEAFY HULL STERILE1* (*LHS1*) with the rOg-PAN-22 region (RL); and *Ghd7.1/Hd2/OsPRR37* with the rOg-PAN-65 (RL) region (Table 2).

### RYMV resistance

Our association study of quantitative RYMV resistance detected a total of 2,199 associated SNPs (Table 1; Additional file 4; Additional file 6: Table S2). These SNPs defined 257 genomic regions containing 2,248 annotated genes (Table 1; Additional file 7: Table S3; Additional file 8: Table S4). The EMMA model revealed a much higher number of significant SNPs compared to the other models, 1,831 SNPs and 177 regions being detected only with this model. Regions that were the most consistent across the different statistical methods were observed on chromosome 3 (rOg-RYMV-125, position 10.5Mb, Additional file 7: Table S3), chromosome 4 (rOg-RYMV-159/rOg-RYMV-160, position 25-25.3Mb), chromosome 6 (rOg-RYMV-200, position 15.1Mb) and chromosome 11 (rOg-RYMV-63, position 9,0-9.2Mb; rOg-RYMV-77/rOg-RYMV-78, position 26.2-26.8Mb).

We detected significant associations in the vicinity of two known resistance genes: *RYMV1*, located in rOg-RYMV-159 on chromosome 4; and *RYMV3*, located in rOg-RYMV-77 on chromosome 11 (Albar et al. 2006; Pidon et al. 2017) (Fig. 2; Table 2). Analysis of GO terms (Additional file 10: Table S6) revealed enrichment in genes from the Golgi apparatus, known to be involved in both the replication of some plant viruses and in intracellular trafficking and cell-to-cell movement (Pitzalis and Heinlein 2018).

**Figure 2.**
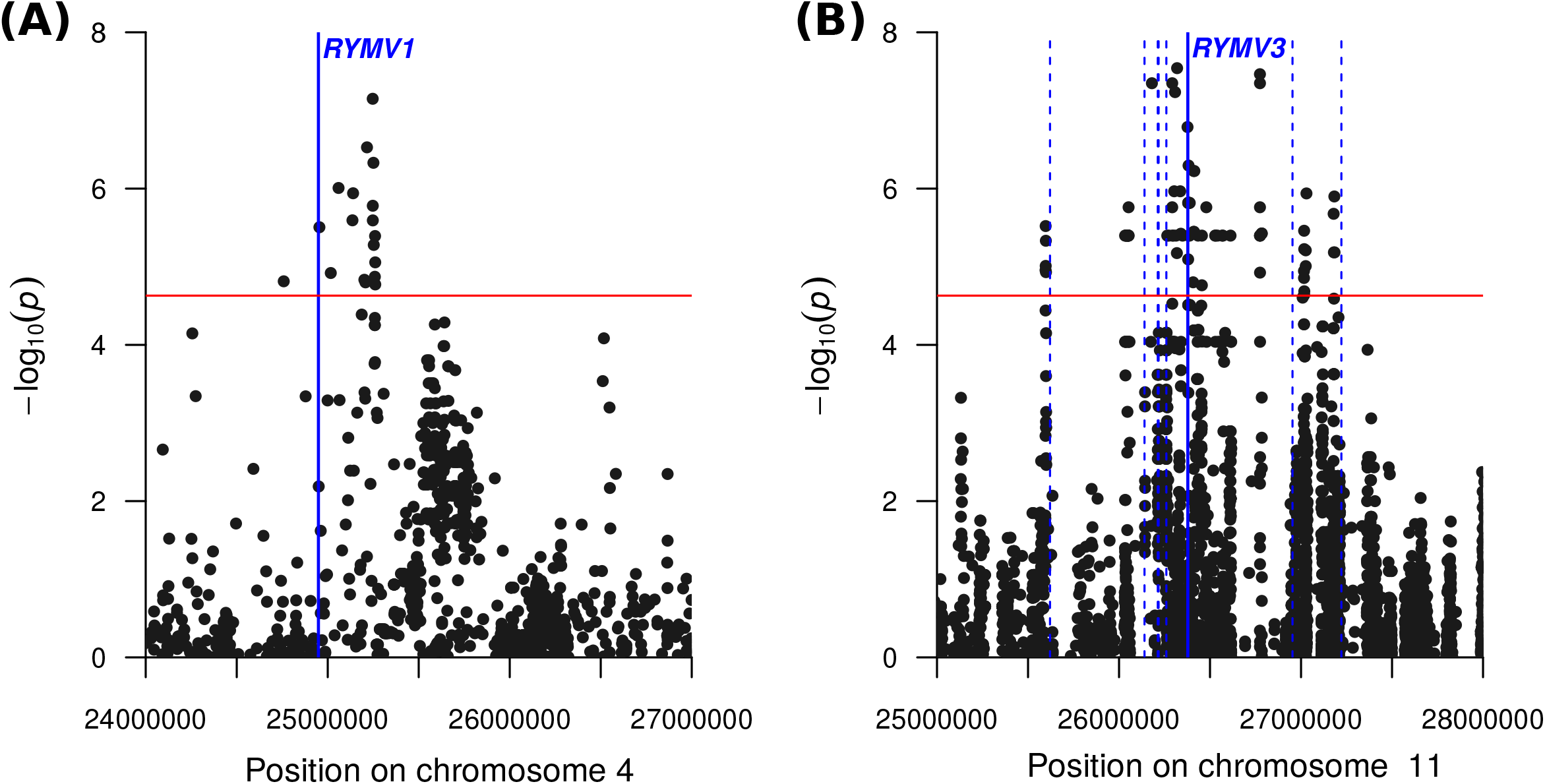
Details of Manhattan plots obtained with LFMM on regions associated to RYMV resistance on chromosome 4 (A) and 11 (B). The 5% FDR threshold is represented by a red horizontal line. Positions of the major resistance genes *RYMV1* and *RYMV3* are indicated by plain blue lines and positions of other *NLR* genes on chromosome 11 are indicated by dotted blue lines.

### Environment-related variables

In order to study association with environmental variables, we downloaded climatic variables for 107 geolocalised accessions (Cubry et al. 2018) from the worldclim v1.4 database (Hijmans et al. 2005). The first two axes of the PCA of bioclimatic data (BioPC1 and BioPC2) explained 49.91% and 26.32% of the variance of all variables. BioPC1 was mainly explained by bio4 (temperature seasonality) and bio17 (precipitation of driest quarter) while BioPC2 was mainly explained by bio9 (mean temperature of driest quarter) and bio11 (mean temperature of coldest quarter). We evaluated the statistical association of genetic polymorphisms with the first and second axes of the PCA. No significant association was detected with BioPC1. BioPC2 allowed the identification of 52 SNPs (Table 1; Additional file 5; Additional file 6: Table S2) and 17 regions encompassing 128 annotated genes (Additional file 7: Table S3; Additional file 8: Table S4). Among the identified genes, we found a cation/proton exchanger-encoding gene (*LOC_Os02g04630, OsCAX4*), an auxin-responsive gene (*LOC_Os03g53150, OsAUX*) and a MADS-box gene (*LOC_Os03g11614, OsMADS1*) (Table 2). GO term enrichment tests detected only one GO term as significantly over-represented, GO:0005576 (Cellular Components / extracellular region) (Additional file 10: Table S5).

When considering maximum temperature variables (Tmax), 73.56% and 17.90% respectively of the variance was explained by the first PCA axis (TmaxPC1) and second PCA axis (TmaxPC2). No association was found with TmaxPC1, while TmaxPC2 allowed us to detect 228 regions including 1,946 annotated genes (Table 1; Additional file 7: Table S3 and Additional file 8: Table S4). GO term analysis revealed several GO terms as being significantly enriched, including seven terms for Biological Processes (cell differentiation, cellular protein modification process, cellular component organization, embryo development, signal transduction, regulation of gene expression and response to external stimulus), two terms for Cellular Components (mitochondrion and plasma membrane) and five terms for Molecular Functions (receptor activity, kinase activity, nucleotide binding, translation factor activity/RNA binding and transferase activity) (Additional file 10: Table S5). One genomic region, rOg-Tmax-36, co-localized with a known QTL for seedling cold tolerance in *O. sativa* (Kim et al. 2014).

## DISCUSSION

### Overlap of flowering time and panicle architecture genetic networks between African and Asian crop species

The flowering pathway is a well described pathway in Asian rice *O. sativa*, with several known key genes (Tsuji et al. 2011; Hori et al. 2016). Based on our expert list of known flowering genes, we were able to show that our method significantly retrieved genes involved in the variability of this trait. We detected candidate regions that co-localize with some previously described flowering genes. Among them, *Hd3a* and *RFT1* are florigen genes homologous to *Arabidopsis thaliana FT* (Komiya et al. 2008). *Hd6* is a flowering repressor, causing late flowering under Long Day (LD) conditions (Ogiso et al. 2010). Overexpression of *OsMADS14*, a homolog *of A. thaliana AP1*, has been shown to be responsible for extreme early flowering in Asian rice (Jeon et al. 2000). *OsMADS51* is a type I MADS-box gene that has been shown to promote flowering in Short Day (SD) conditions (Kim et al. 2007). *OsGI* is an ortholog of the *A. thaliana GIGANTEA* gene and its over-expression in rice leads to late flowering under both SD and LD conditions (Hayama et al. 2003). *ETR2*, encoding an ethylene receptor, has been shown to delay flowering by regulating the expression of *OsGI* (Wuriyanghan et al. 2009; Tsuji et al. 2011). Functional characterization of the *RCN1* gene, homologous to *A. thaliana TFL1*, showed that this gene can promote late-flowering when overexpressed in transgenic plants (Nakagawa et al. 2002). Finally the phytochrome *OsPhyC* gene encodes one of the three rice phytochromes and has been shown to be a flowering repressor (Takano et al. 2005).

A large number of additional genomic regions (Additional file 7: Table S3) were identified for which further studies should be conducted in order to identify novel genetic diversity relating to the flowering pathway of African rice. The large number of regions observed is to be expected when dealing with polygenic traits and it should be noted that a comparable number of low effect QTLs was previously reported for flowering time in Asian rice on the basis of genetic mapping studies (Hori et al. 2016).

Spikelet number per panicle was the main trait contributing to the diversity of panicle architecture observed in this population. No significant SNPs were identified for the SpN trait using any of the 4 models. A possible explanation for this result is that the trait might be associated with a large number of QTLs of low effect sizes, and may consequently be difficult to assess using the present GWAS panel. For panicle traits, the detected peaks were broadly characterized by relatively high *p*-values, suggesting that the associated morphological traits are highly polygenic with small effect sizes. Some genes previously implicated in the regulation of panicle development and/or architecture in *O. sativa* (Wang and Li 2011; Teo et al. 2014) were also found to be associated with panicle morphological trait variations in *O. glaberrima*, indicating a parallel evolution of the trait in the two species. The *APO1* gene encodes an ortholog of the *A. thaliana* UNUSUAL FLOWER ORGAN (UFO) F-box protein and was reported to be involved in the control of rice spikelet number and panicle branching complexity through effects on meristem fate and cell proliferation (Ikeda et al. 2007; Ikeda-Kawakatsu et al. 2009). This gene was also found to be associated with QTLs relating to yield, panicle architecture complexity and lodging resistance in *O. sativa* (Terao et al. 2010; Ookawa et al. 2010). The *OsMADS1*/*LEAFY HULL STERILE1* (*LHS1*) gene encodes a *SEPALLATA*-like MADS-box transcription factor (TF), which promotes the formation of spikelet/floret meristems (Jeon et al. 2000; Agrawal et al. 2005; Khanday et al. 2013). Interestingly, this gene was also identified within a major drought-tolerant QTL (qDTY3.2) (Saikumar et al. 2014) from a cross between *O. sativa* and *O. glaberrima*, corresponding to a genomic region associated with response to environmental variables (rOg-BioClim-3). *SHORT PANICLE1* (*SP1*) encodes a putative PTR family transporter, which controls panicle size and branching complexity (Li et al. 2009). The *Ghd7.1/Hd2/OsPRR37* gene encodes a *PSEUDO-RESPONSE REGULATOR* (*PRR*) protein associated with a QTL displaying pleiotropic effects on spikelet number per panicle, plant height and heading date (Liu et al. 2013).

Several association studies of panicle morphological trait diversity have been recently conducted for *O. sativa* (Bai et al. 2016; Crowell et al. 2016; Rebolledo et al. 2016; Ta et al. 2018; Yano et al. 2019). Only a few overlaps of GWAS candidates were observed between the two rice crop species. As for the flowering time analysis, a large number of specific genomic regions (Additional file 7: Table S3) were identified. For these regions, further studies should lead to the precise identification of genetic elements governing panicle diversity in African rice. Among the genes located in these specific regions, some were previously functionally characterized in *O. sativa* but without any observations of a direct relationship with panicle development, examples including: *OsIPT2* (*ADENOSINE PHOSPHATE ISOPENTENYLTRANSFERASE*), encoding a cytokinin biosynthetic gene regulated by *OsARID3* which is essential for shoot apical meristem (SAM) development in rice (Xu et al. 2015); *VERNALIZATION INSENSITIVE 3-LIKE 1* (*OsVIL1*) which is involved in flowering time through *OsLF* and *GhD7* genes (Jeong et al. 2016); and the *ent*-kaurene synthase genes *OsKS1* and *OsKS2*, which are involved in GA biosynthesis and associated with a GWAS mega locus related to panicle and yield traits in *O. sativa* (Sakamoto and Matsuoka 2004; Crowell et al. 2016). Other genes may be of interest on the basis of their annotations and the known functions of their orthologs, examples including: *OsGRF5* and *GRF7*, two members of the *GROWTH-REGULATING FACTOR* or GRF small gene family encoding TFs associated with SAM maintenance and flowering time in rice (Omidbakhshfard et al. 2015); and *AP2_EREBP77* and *OsRAV2*, two members of the *RELATED TO ABI3 AND VP1* or *RAV* subfamily of AP2/ERF TFs, homologous to the *TEM* genes from *A. thaliana* that affect FT induction through the photoperiod and GA pathways (Swaminathan et al. 2008).

### Quantitative resistance to RYMV in *O. glaberrima* and major resistance genes

The large number of significant SNPs associated with RYMV resistance may reflect the highly quantitative nature of partial resistance. However 83% of significant SNPs were detected only by the EMMA method (1,831 SNPs), suggesting that a high number of false positive SNPs was detected with this method.

The regions identified as being associated with resistance against RYMV did not overlap with QTLs of partial resistance against RYMV previously identified in *O. sativa* (Boisnard et al. 2007), suggesting that different genes and pathways may lead to resistance. However, two major resistance genes against RYMV, *RYMV1* and *RYMV3*, were found to be associated with quantitative resistance. *RYMV1* encodes a translation initiation factor that acts as a susceptibility factor through its interaction with the genome-linked protein (VPg) of RYMV, which is required for viral infection (Albar et al. 2006; Hébrard et al. 2010). Several different alleles conferring high resistance are known (Thiémélé et al. 2010). The main candidate resistance gene for *RYMV3* belongs to the family of nucleotide-binding domain and leucine-rich repeat containing (NLR) genes (Pidon et al. 2017), many of which are involved in pathogen recognition and effector-triggered immunity (de Ronde et al. 2014). NLR genes are known to be frequently organized into clusters and several additional NLR genes, annotated in the close vicinity of the *RYMV3* candidate gene, might also be good candidates for quantitative resistance. Translation initiation factors and NLRs are known to act as determinants of high and monogenic resistance (de Ronde et al. 2014; Sanfaçon 2015). Moreover, a role in quantitative resistance has been clearly established for NLRs (Wang et al. 1999; Hayashi et al. 2010) and suggested for translation initiation factors (Nicaise et al. 2003; Marandel et al. 2009). The *RYMV1* and *RYMV3* loci, or adjacent NLR gene loci, might thus harbor both alleles with quantitative effects and alleles with strong effects on RYMV resistance. Similarly, even if the *RYMV2* major resistance gene to RYMV was not found to be associated with partial resistance in this study, a previous analysis strongly suggested that *RYMV2* might be involved in partial resistance to RYMV in the species *O. sativa* (Orjuela et al. 2013).

In addition to the above, genes encoding protein domains implicated previously in virus resistance, such as lectin (Yamaji et al. 2012) or methrin-TRAF domains (Cosson et al. 2010), were also found to be located in or close to significantly associated regions (Additional file 8: Table S4) and may constitute interesting candidates. Further studies should be conducted on these candidate genes, as well as on the additional genomic regions identified (see Additional file 7: Table S3) in order to describe the diversity of resistance pathways to RYMV in African rice.

### Relationship between environment-related variables and *O. glaberrima* diversity

We did not find any significant association between the first PCA axis and either the set of bioclimatic variables or the monthly average maximal temperature. This could be explained by a high correlation between the first PCA axis and population genetic structure (Frichot et al. 2015). Correcting for confounding effects did not allow us to detect association at SNPs displaying allelic frequencies that correlated with such a structure. For the second PCA axis, several significant associations were found and interesting genes were identified, including genes related to drought or cold tolerance. Among the most interesting candidates, *OsCAX4* is a salt-tolerance gene (Molla et al. 2015), *OsAUX* is an auxin-responsive gene that has been found to be significantly over-expressed in cold-tolerant seedlings (Dametto et al. 2015) and *OsMADS1* is located within a major drought-tolerant QTL (qDTY3.2) detected from a cross between both *O. sativa* and *O. glaberrima* (Saikumar et al. 2014).

Several candidate regions have been identified that will require a more in-depth study in order to gather variations of interest for breeding purposes (see Additional file 7: Table S3). This is especially true for the temperature-related candidates, the latter being likely targets for genetic improvement in the context of a changing climate.

## CONCLUSIONS

We report on the results of an extensive Genome Wide Association Study carried out for several traits on African rice. Analysis of the well-studied character of flowering time enabled us to retrieve a significant number of previously identified genes, thus validating our approach. We also carried out the first GWAS analysis to date of climate variables in relation to African rice, obtaining a list of candidate regions that should be the subject of further studies in order to detect functional variation linked to local adaptation and resistance or tolerance to abiotic stresses.

The RYMV resistance and panicle architecture traits for which we report significant associations are key agronomic traits that should interest farmers and breeders. Further studies aiming at identifying adaptive polymorphisms among the candidates found in this study and functional validation of candidate genes will be needed to reinforce our results.

## MATERIAL AND METHODS

### Genotypic data

Single nucleotide polymorphisms (SNPs) from 163 high-depth re-sequenced *O. glaberrima* accessions were used in this study (Cubry et al. 2018). SNPs were identified based on mapping to the *Oryza sativa japonica* cv. Nipponbare high quality reference genome in terms of assembly and annotation (Kawahara et al. 2013). The bioinformatic mapping pipeline, software and SNP filtering steps that were used are described in Cubry et al. (2018).

SNPs with more than 5% missing data (minor fraction of total SNP set) were filtered out (Cubry et al. 2018). As missing data can reduce the power of association studies (Browning 2008; Marchini and Howie 2010), we imputed the remaining missing data based on a matrix factorization approach using the “impute” function from the R package LEA (Frichot and François 2015). This approach uses the results f ancestry estimation from a sparse non-negative matrix factorization (sNMF) analysis to infer missing genotypes (Frichot et al. 2014). In sNMF, we set K to infer four clusters and kept the best out of 10 runs based on a cross entropy criterion.

### Phenotyping of flowering time and panicle morphology

Phenotyping of flowering time and panicle morphology was performed near Banfora (Burkina-Faso) under irrigated field conditions at the Institut de l’Environnement et de Recherches Agricoles (INERA) station in 2012 and 2014. Plants were sown at two different periods in the same year: the first at beginning of June (“early sowing”) and second in mid-July (“late sowing”). A total of 15 plants per plot of 0.5 m^2^ were grown. The field trials followed an alpha-lattice design with two replicates (Patterson and Williams 1976) per date of sowing per year. Each single block included 19 accessions (i.e. 19 plots). In total, 87 *O. glaberrima* accessions were planted in 2012 and 155 in 2014.

Flowering date (DFT) was scored when 50% of the plants for a given accession harbored heading panicles. Fourteen days after heading date, the three main panicles from three central plants per plot per repeat were collected (i.e. nine panicles/accession/repeat) from the early sowing. Each panicle was fixed on a white paper board, photographed and phenotyped using the P-TRAP software (AL-Tam et al. 2013). We estimated rachis length (RL), primary branch number (PBN), primary branch average length (PBL), primary branch internode average length (PBintL), secondary branch number (SBN), secondary branch internode average length (SBintL) and spikelet number (SpN). All statistical analyses of the dataset were performed using R packages ade4, corrplot and agricolae (R software version 1.2.1335) as described in Ta et al. (2018).

### RYMV resistance phenotyping

Resistance was evaluated based on ELISA performed on infected plants cultivated in the greenhouse, under controlled conditions. As high resistance to RYMV is widely accepted for African rice, we excluded highly resistant accessions, i.e. in which no virus can be detected with ELISA (Thiémélé et al. 2010; Orjuela et al. 2013; Pidon et al. 2017), and we focused only on quantitative resistance. We therefore assessed resistance on a set of 125 accessions. Two varieties were used as susceptibility controls, IR64 (*O. sativa ssp. indica*) and Nipponbare (*O. sativa ssp. japonica*), and one as a high resistance control, Tog5681 (*O. glaberrima*). Three replicate experiments of all varieties were performed. In each experiment, plants were organized in two complete blocks with four plant replicates per accession.

Plants were mechanically inoculated three weeks after sowing, as described in Pinel-Galzi et al. (2018) with CI4 isolate of RYMV (Pinel et al. 2000). Four discs of 4 mm diameter were cut on the last emerged leaf of each plant 17 and 20 days after inoculation (dai) and discs from the four plants of the same repeat were pooled. Samples were ground with a QIAGEN TissueLyser II bead mill and resuspended in 750 µL 1X PBST (Phosphate buffer saline with Tween 20).

Virus content was estimated by DAS-ELISA (Pinel-Galzi et al. 2018). Preliminary tests on a subset of samples were performed to assess the dilution that best discriminated between samples. ELISA tests were finally performed at dilutions of 1/1,000 for 17 dai sampling date and 1/2,500 for 20 dai sampling date. As virus content was highly correlated between 17 and 20 days after infection (R^2^=0,81), the resistance level was estimated as the mean of the two sampling dates.

### Environmental variables

For accessions with geographical sampling coordinates, we retrieved information for 19 climate-related variables (referred to here as bio) from the WORLDCLIM database at a 2.5 minute resolution (Hijmans et al. 2005). We also retrieved the average monthly maximum temperature (referred to here as Tmax). We first performed a Principal Component Analysis (PCA) on each set of variables to build uncorrelated composite variables. PCA were performed using R software (Frichot and François 2015). Association studies were performed using the first two components of each PCA.

### Association studies

For each trial, SNPs displaying a minimal allele frequency (frequency of the minor allele) lower than 5% were filtered out. We first adjusted a simple linear model (Analysis of variance, ANOVA) to associate phenotype and genotype. This simple method did not take into account any putative confounding factor and allowed us to assess whether taking into account relatedness and/or population structure could reduce false positive rates. Two classes of methods accounting for confounding factors were used: 1) mixed models using kinship matrix and/or population structure (Yu et al. 2006); and 2) latent factor methods (Frichot et al. 2013). We used both mixed linear models MLM (Zhang et al. 2010) as implemented in GAPIT R package (Lipka et al. 2012) and EMMA (Kang et al. 2008) as implemented in R package EMMA. For EMMA, the kinship matrix was estimated using the emma.kinship function. For MLM (Q+K model), the kinship (K matrix) was computed using the Van Raden method and the first three principal components of a PCA of genomic data were used as the Q matrix. Finally, we used latent factor methods (Frichot et al. 2013) that jointly estimated associations between genotype and phenotype and confounding factors. We used the R packages LFMM2 (Caye et al. 2019) and CATE (Wang et al. 2017) to perform these analyses. For LFMM2, we first made the estimation of the confounding factors by using a subset of SNPs obtained by applying a 20% MAF filter, and we considered four latent factors (Cubry et al. 2018). We then used the resulting confounding matrix for the analysis of genotype/phenotype association. For CATE, we considered all SNPs and we assumed four confounding factors in the association model. The results of all analyses were graphically represented by using a QQ-plot to assess confounding factor correction and Manhattan plots (R package qqman, Turner 2014). We used a false discovery rate (FDR) of 5% to select candidate SNPs for each method. FDR estimation was realized using the R package qvalue (Storey et al. 2019).

GWAS analysis was performed separately for each year and trial (see Additional file 1: Table S1). *P*-values obtained for the same traits or the same planting data were combined across experiments using Fisher’s method (Sokal and Rohlf 2012). The final list of candidate SNPs was established for each trait by considering SNPs detected by at least one method. Annotation of retained candidate SNPs was performed using the SNPeff annotation data for MSU7 (Kawahara et al. 2013), considering genes within the region 25kb upstream and 25kb downstream from each detected SNP. We used the funRiceGenes database (http://funricegenes.ncpgr.cn; Yao et al. 2018) to extract gene symbols and associated functional information whenever possible.

Finally, for flowering traits, we established a list of known genes of particular interest from published data (Tsuji et al. 2011; Hori et al. 2016). This “expert” list was then used to assess the performance of our GWAS approach to retrieve these potential candidates. We used a G-test to assess enrichment of candidates in our list of identified genes.

Gene ontology (GO) term enrichment tests for biological process, cellular component and molecular function terms were performed using the Fisher exact test implemented in the R package TopGO (Alexa et al. 2006).

## List of abbreviations

ANOVA: Analysis of Variance
BioPC1: First principal component of PCA on bioclimatic variables
BioPC2: Second principal component of PCA on bioclimatic variables
dai: Days after inoculation
DFT: Days to flowering
ELISA: Enzyme-linked immunosorbent assay
FDR: False discovery rate
GO: Gene ontology
GWAS: Genome wide association study
INERA: Institut de l’environnement et de recherches agricoles
LD: Long days
NLR: Nucleotide binding leucine rich repeat
PbintL: Primary branch internode average length
PBL: Primary branch average length
PBN: Primary branch number
PCA: Principle component analysis
QTL: Quantitative trait locus
QQ-plot: quantile-quantile plot
RL: Rachis length
RYMV: Rice yellow mottle virus
SbintL: Secondary branch internode average length
SBN: Secondary branch number
SD: Short days
SpN: Spikelet number
SNP: Single nucleotide polymorphism
TF: Transcription factor
TmaxPC1: First principal component of PCA on maximal temperature variables
TmaxPC2: Second principal component of PCA on maximal temperature variables

## Declarations

### Ethics approval and consent to participate

Not applicable

### Consent for publication

Not applicable

### Availability of data and material

All customized R scripts are available as a GitHub repository: https://github.com/Africrop/gwas_african_rice. The repository also contains the imputed genotypic data used here to reproduce exactly the same analysis. Phenotypic data are provided as supplemental material.

### Authors’ contributions

YV, LA, SJ and FS planned and supervised the study. HP, TKN, SJ, LA, HA, HK, HC and AG participated in data generation and collection management. PC, TKN, HP, CTD, ACT, MH, OF, FS, YV, SJ and LA participated in the statistical analysis. PC, HP, FS, YV, LA and SJ wrote the manuscript. All the authors read and agreed the manuscript.

## Acknowledgements

We thank Ndomassi Tando and the IRD itrop “Plantes Santé” bioinformatic platform for providing HPC resources and support for our research project. We thank Mikael Valéro for technical help in RYMV resistance evaluation. The authors thank staff from INERA station at Banfora (Burkina Faso) for support for field experiment and phenotype scoring, and Ha Thi Loan from LMI RICE (Vietnam) for panicle trait scoring. We also thank Dr James Tregear, IRD, for his careful reading and language editing of the manuscript.

## ADDITIONAL FILE CAPTIONS

**Additional file 1_Table S1. Phenotypic data used for genome-wide association analyses.** Flowering time (DFT), rachis length (RL), primary branch number (PBN), primary branch average length (PBL), primary branch internode average length (PBintL), secondary branch number (SBN), secondary branch internode average length (SBintL) and spikelet number (SpN) were evaluated in field conditions in 2012 and 2014. Resistance to RYMV was evaluated in greenhouse conditions during three experiments (RYMV1, RYMV2, RYMV3) shifted of about 1-2 months. Environmental data were extracted from the worldclim database at available sampling locations. A Principal Component Analysis was then performed on i) the whole set of variables and ii) only maximal temperature related ones. The two first axes of both these PCA were then used for association analyses and are reported in this table.

**Additional file 2. Manhattan plots and QQ plots for flowering time.** Association analysis was performed on both early and late plant sowings. The 5% FDR thresholds are indicated by red lines.

**Additional file 3. Manhattan plots and QQ plots for panicle architecture related traits.** Association analysis was performed independently for each trait and repetition. *P*-values obtained for each replicate were then combined using a Fisher method to obtain final *p*-values for each trait. The 5% FDR thresholds are indicated by red lines.

**Additional file 4. Manhattan plots and QQ plots for RYMV resistance.** The three replicates of phenotypic evaluation were combined based on a Fisher method. The 5% FDR thresholds are indicated by red lines.

**Additional file 5. Manhattan plots and QQ plots for environmental data.** The 5% FDR thresholds are indicated by red lines.

**Additional file 6_Table S2. List of the SNPs associated with eleven different phenotypic traits.** Significant SNPs were identified based on four different models (EMMA, CATE, LFMM and GAPIT), the Fisher method to combine several repetitions of phenotypic data and a 5% FDR threshold. For each trait, the *p*-values obtained with the different models are indicated if significant.

**Additional file 7_Table S3. List of the genomic regions associated with five different categories of phenotypic traits.** Regions were defined based on 50 kb windows around the significant SNPs detected with any of the four models. Overlapping regions were combined into a single one. The chromosome (Chr), the starting (Position 1) and ending (Position 2) positions, the size of the region (Intervals) in base pairs and the number of significant SNPs included (Sign_SNPs_nb) are indicated. Sheets ”RYMV”, “Tmax”, BioClim” concerned the regions identified for the resistance to RYMV, the maximum temperature related variables and the whole set of bioclimatic variables, respectively. The sheet “Flowering” concerned regions identified with the early sowing flowering time (Early) or the late sowing flowering time (Late) traits, as indicated in the two last columns. The sheet “Panicle” concerned regions identified with rachis length (RL), primary branch number (PBN), primary branch average length (PBL), primary branch internode average length (PBintL), secondary branch number (SBN), secondary branch internode average length (SBintL) and spikelet number (SpN), as indicated in the last columns.

**Additional file 8_Table S4. List of genes located in each of the regions associated with five categories of phenotypic traits.**

**Additional file 9_Table S5. Expert list of genes previously described as involved in flowering time in Asian rice and test of enrichment of the list of genes detected by our association analysis.** Fold-enrichment and G-test associated *p*-value are reported.

**Additional file 10_Table S6. GO terms tables.** GO term analyses were run independently for each trait (TmaxPC2, BioPC2, Flowering Time, Panicle, and RYMV). Results are shown for each trait and for the three GO term categories (Biological processes, Cellular components, Molecular functions).

