## Supplementary figures and images for "Genome wide association study pinpoints key agronomic QTLs in African rice *Oryza glaberrima*"

### Additional File 2

#### Early sowing

**Gapit**

**EMMA**

**CATE**

**LFMM**

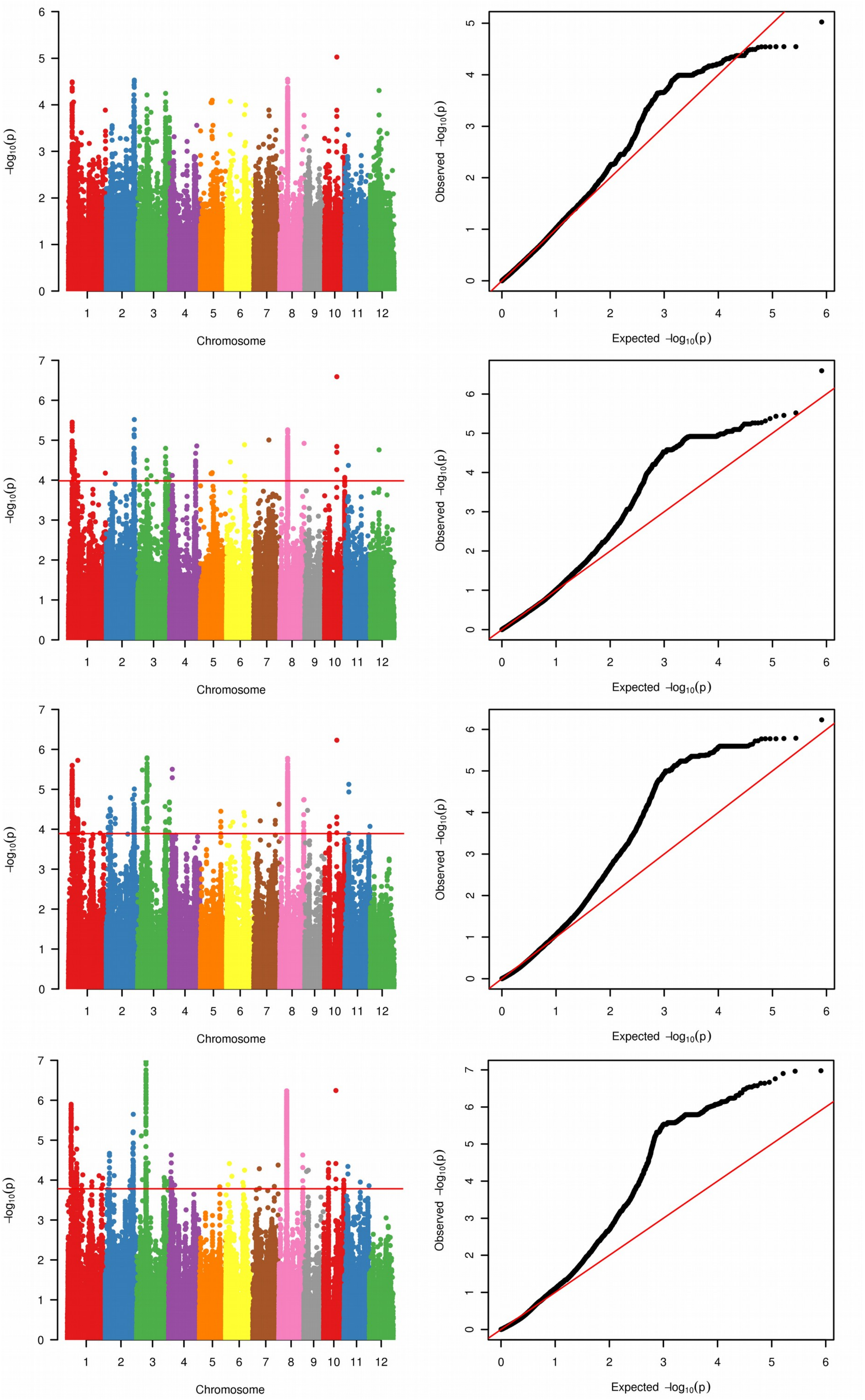

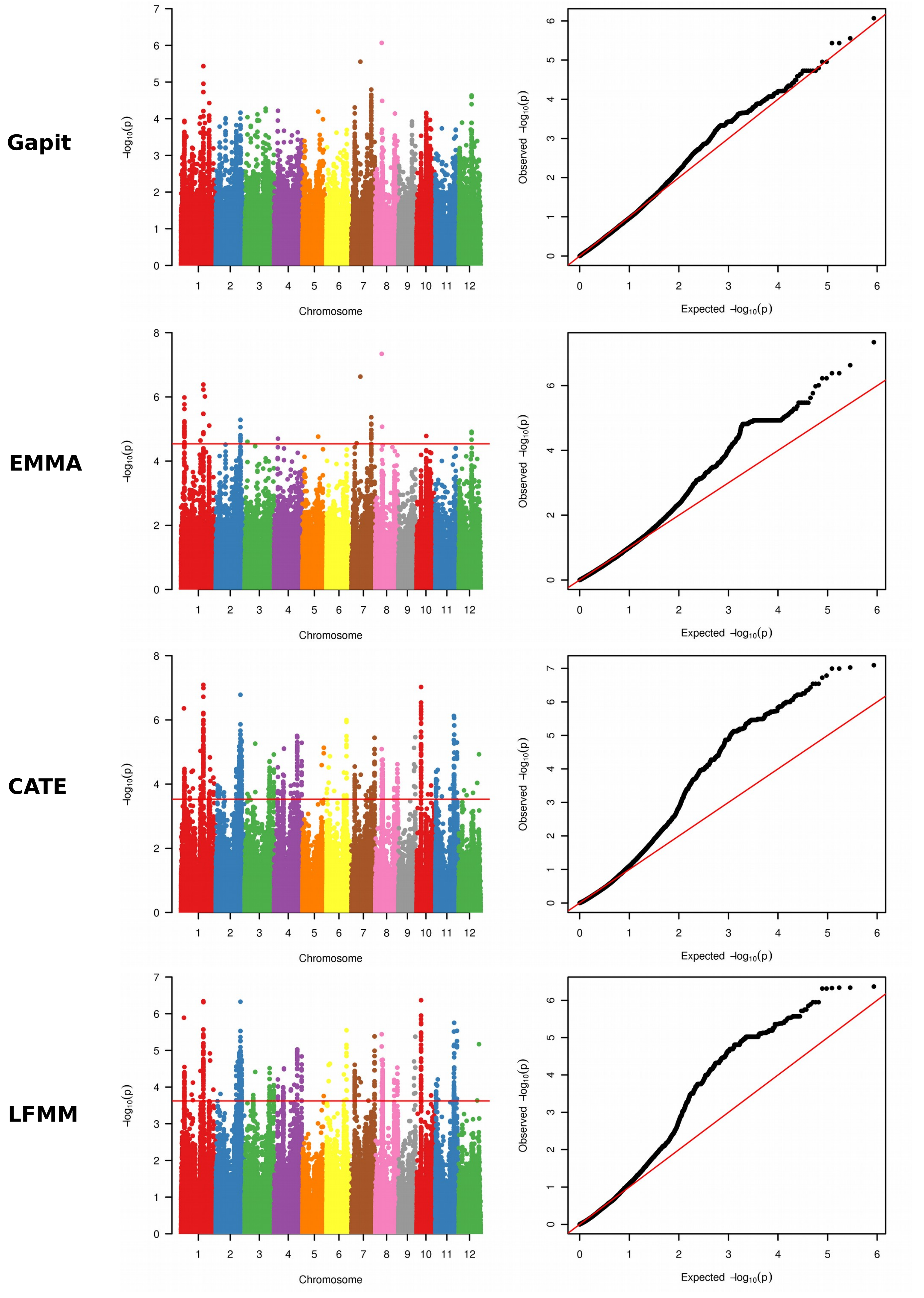

### Additional File 3

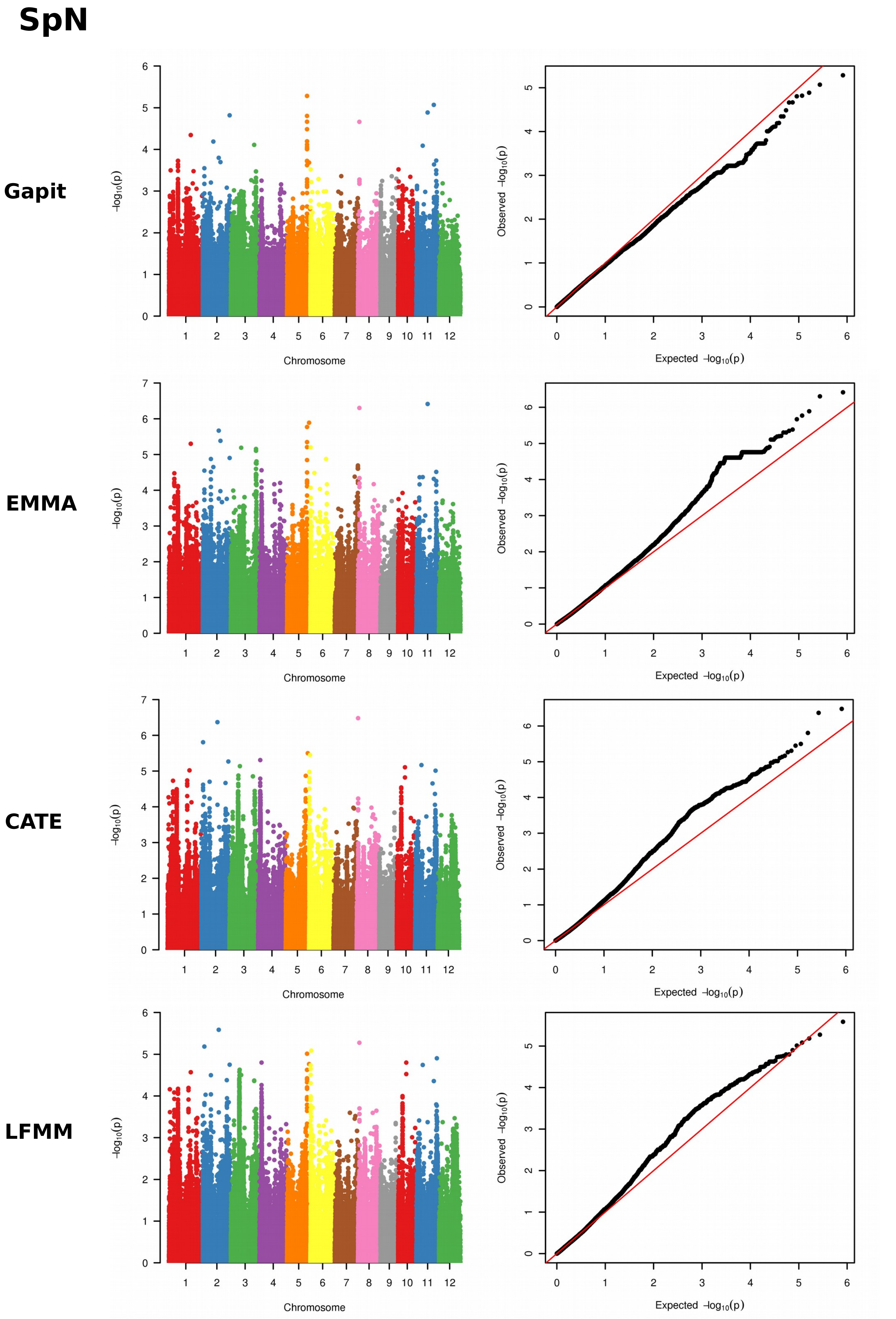

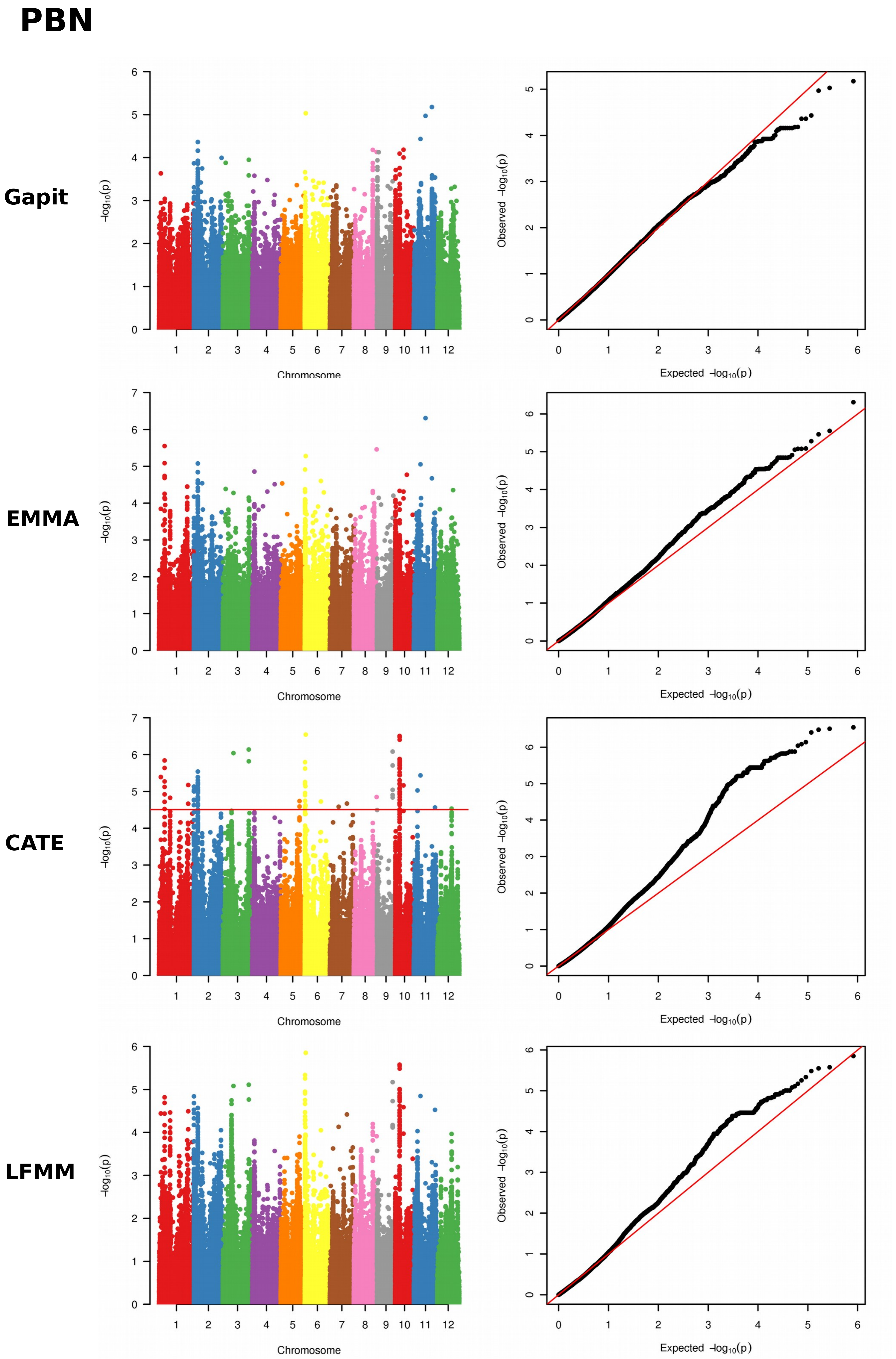

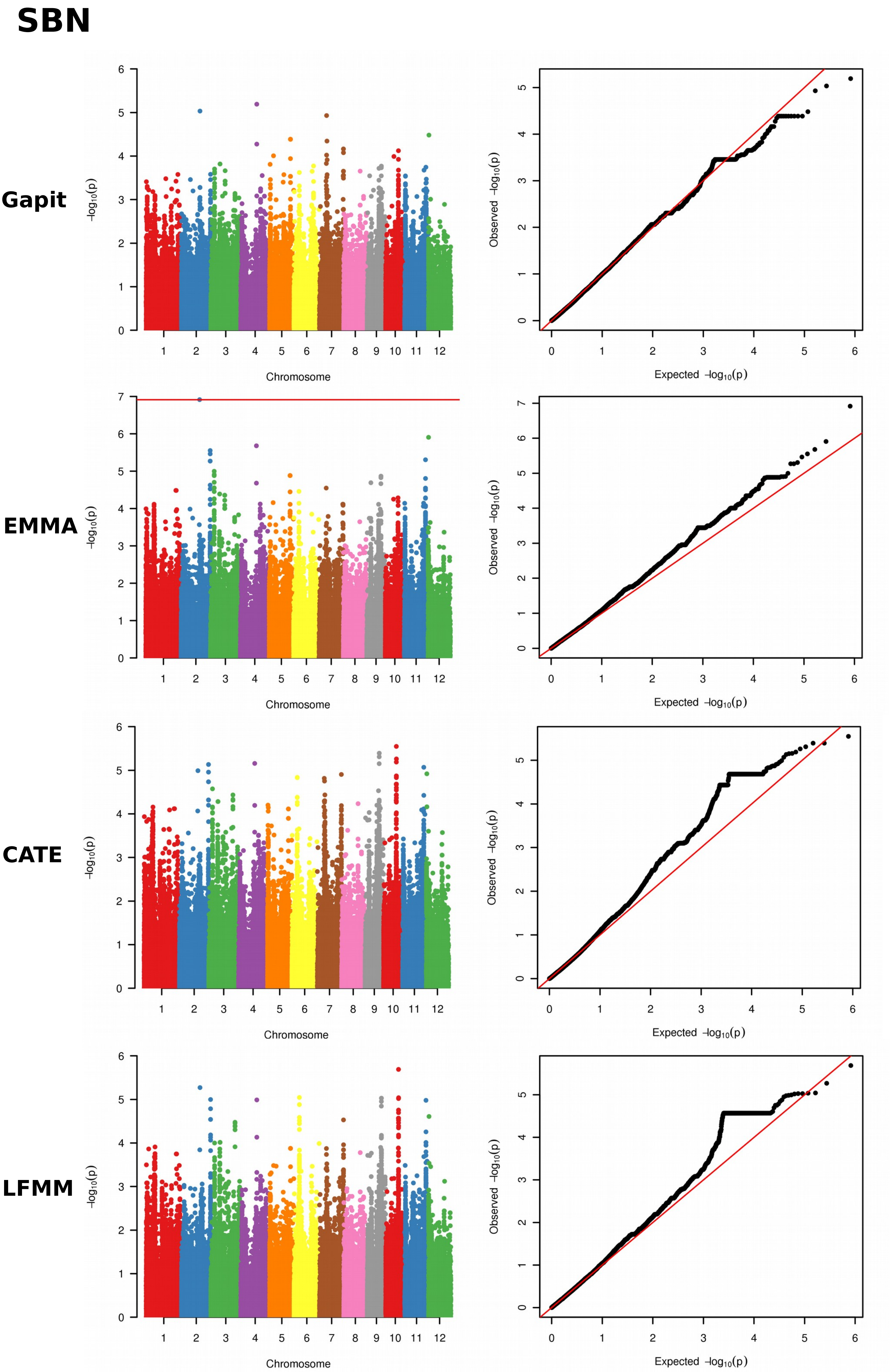

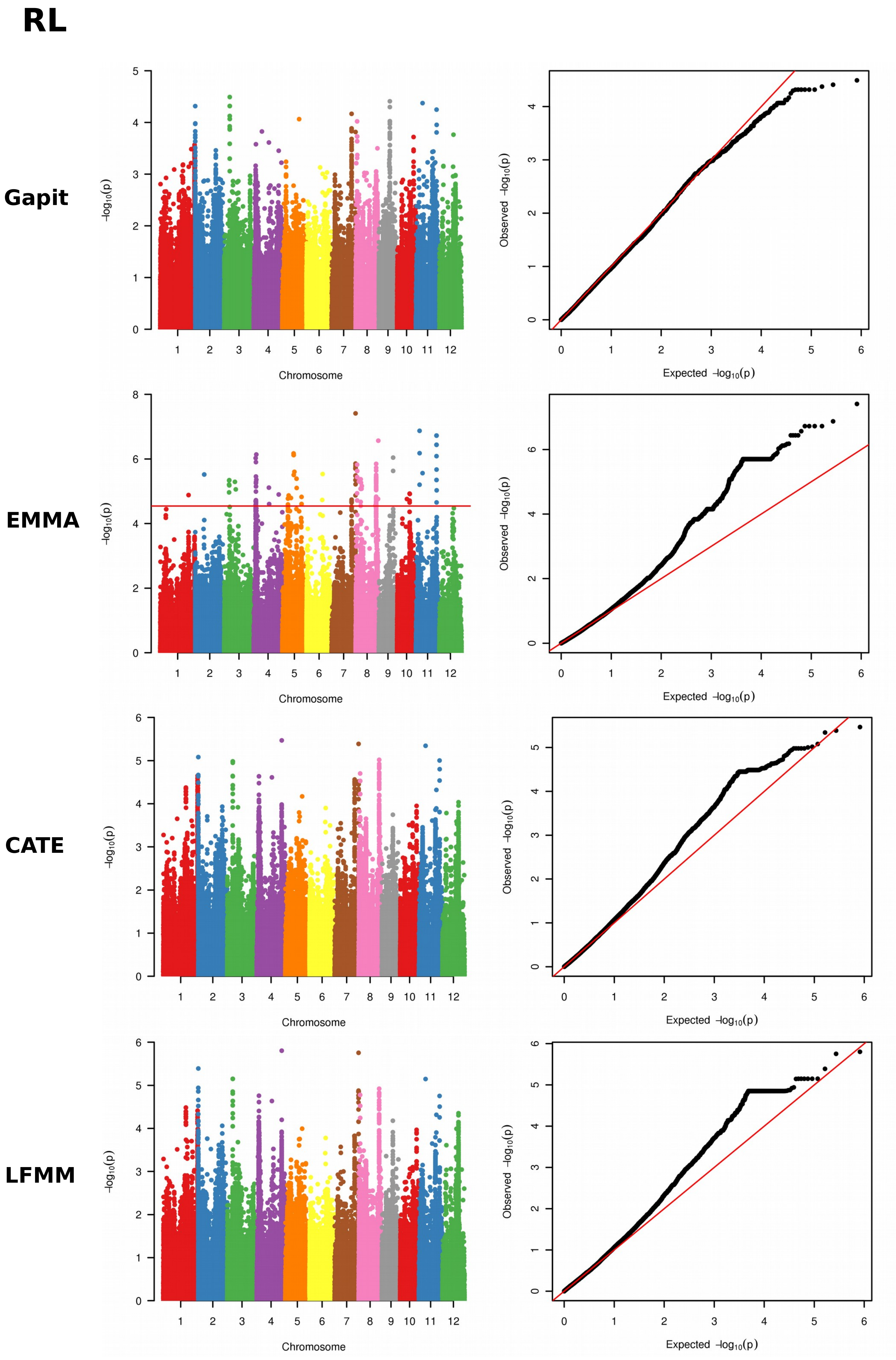

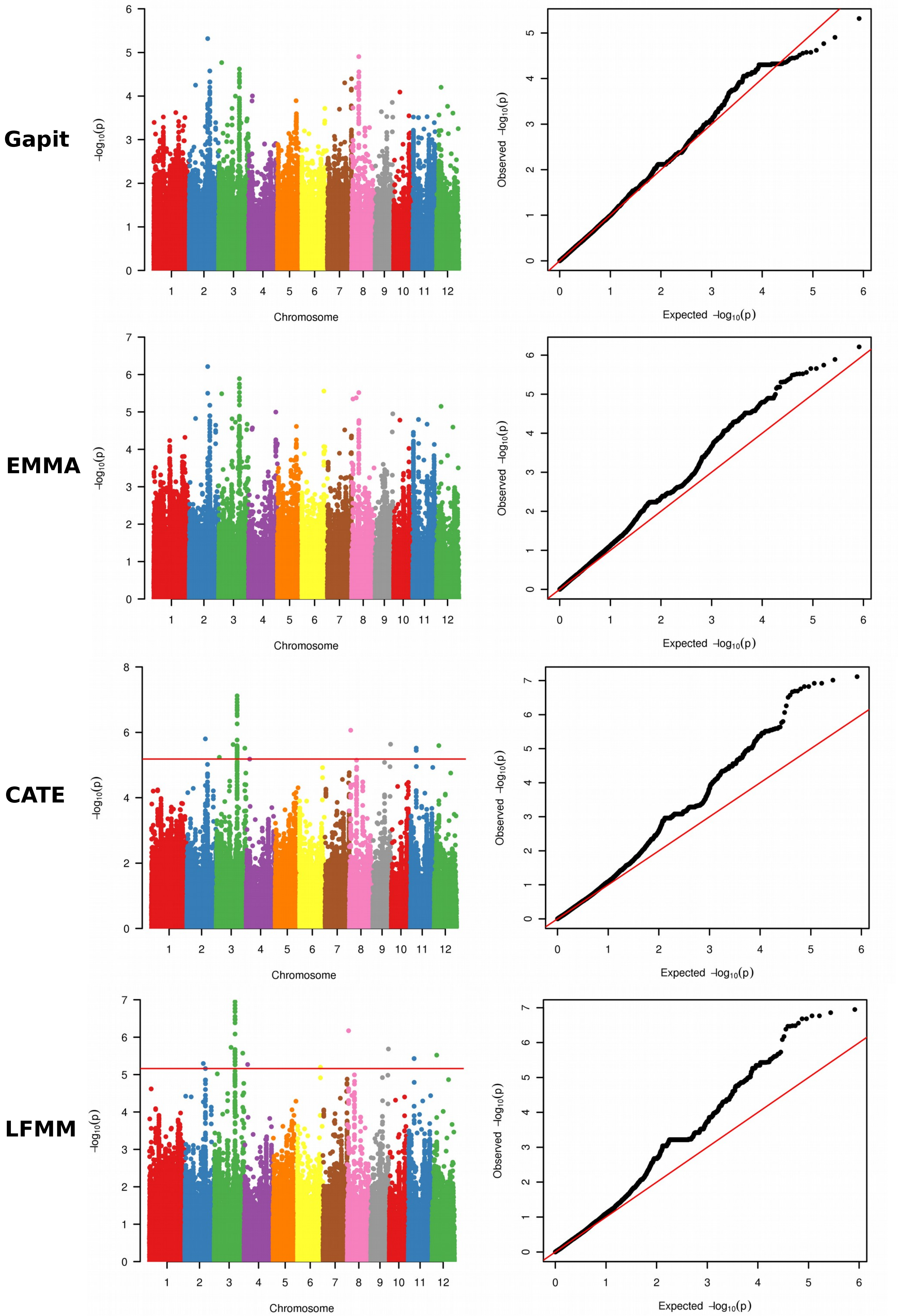

#### PBintL

**Gapit**

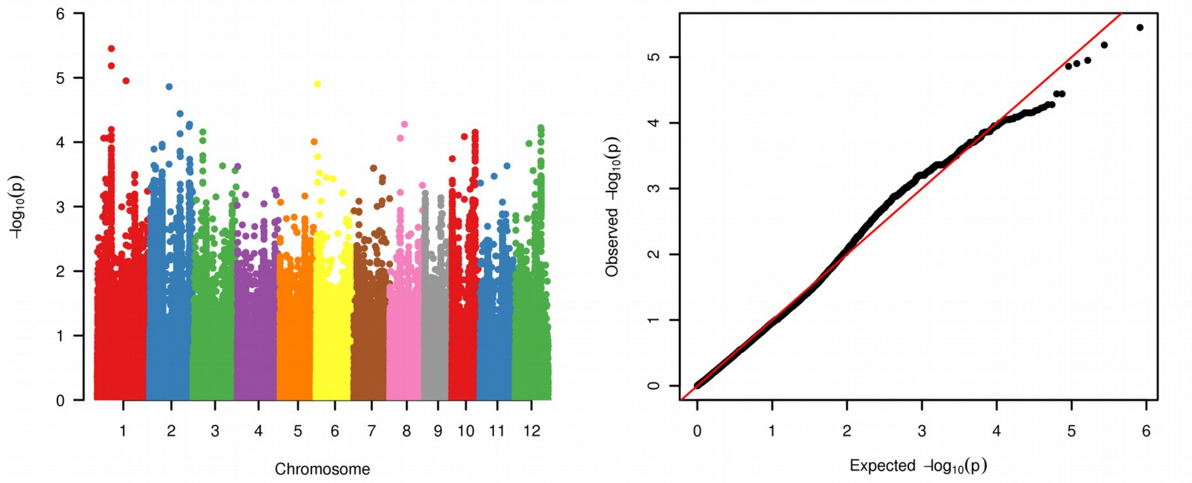


**EMMA**

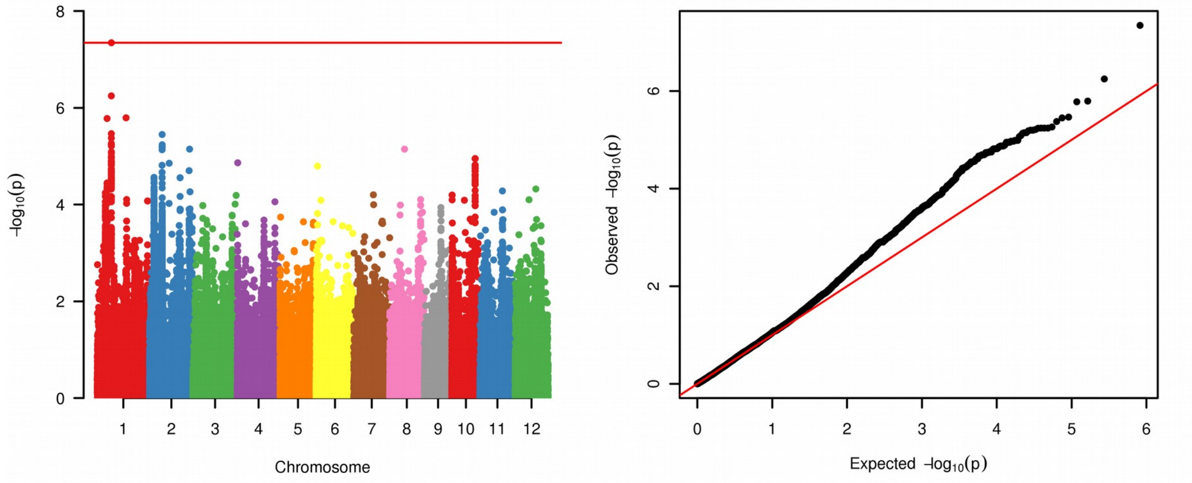


**CATE**

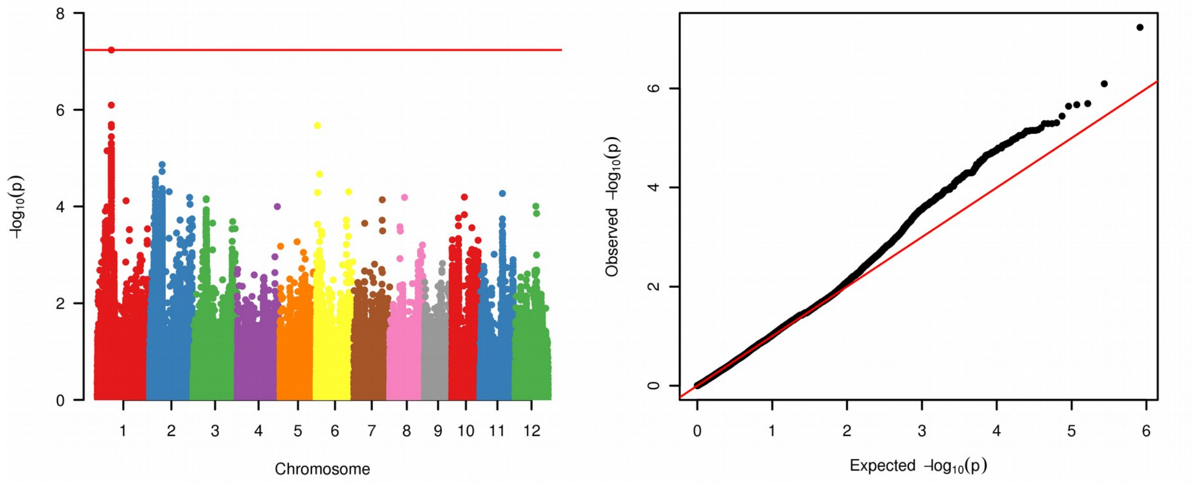


**LFMM**

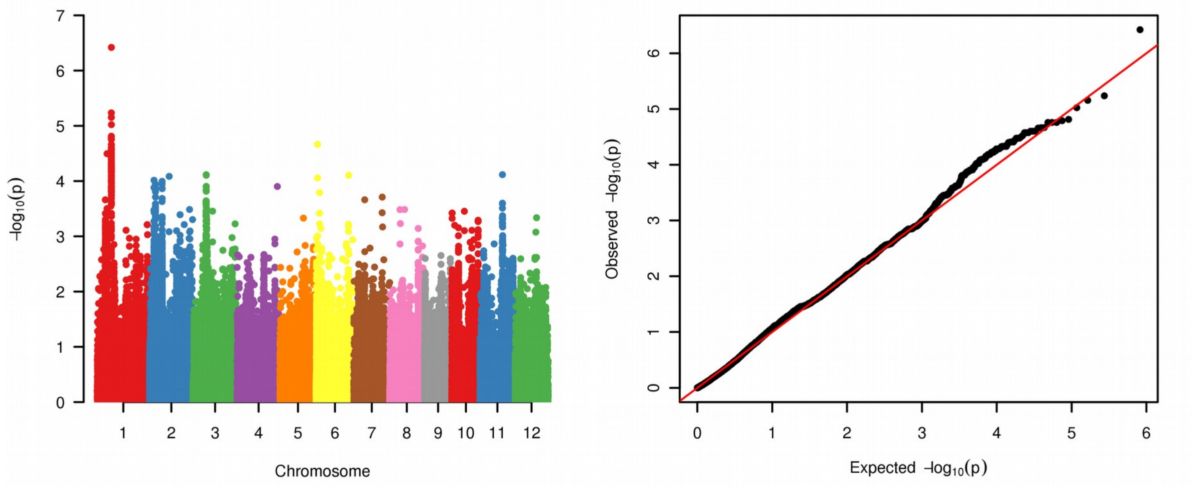

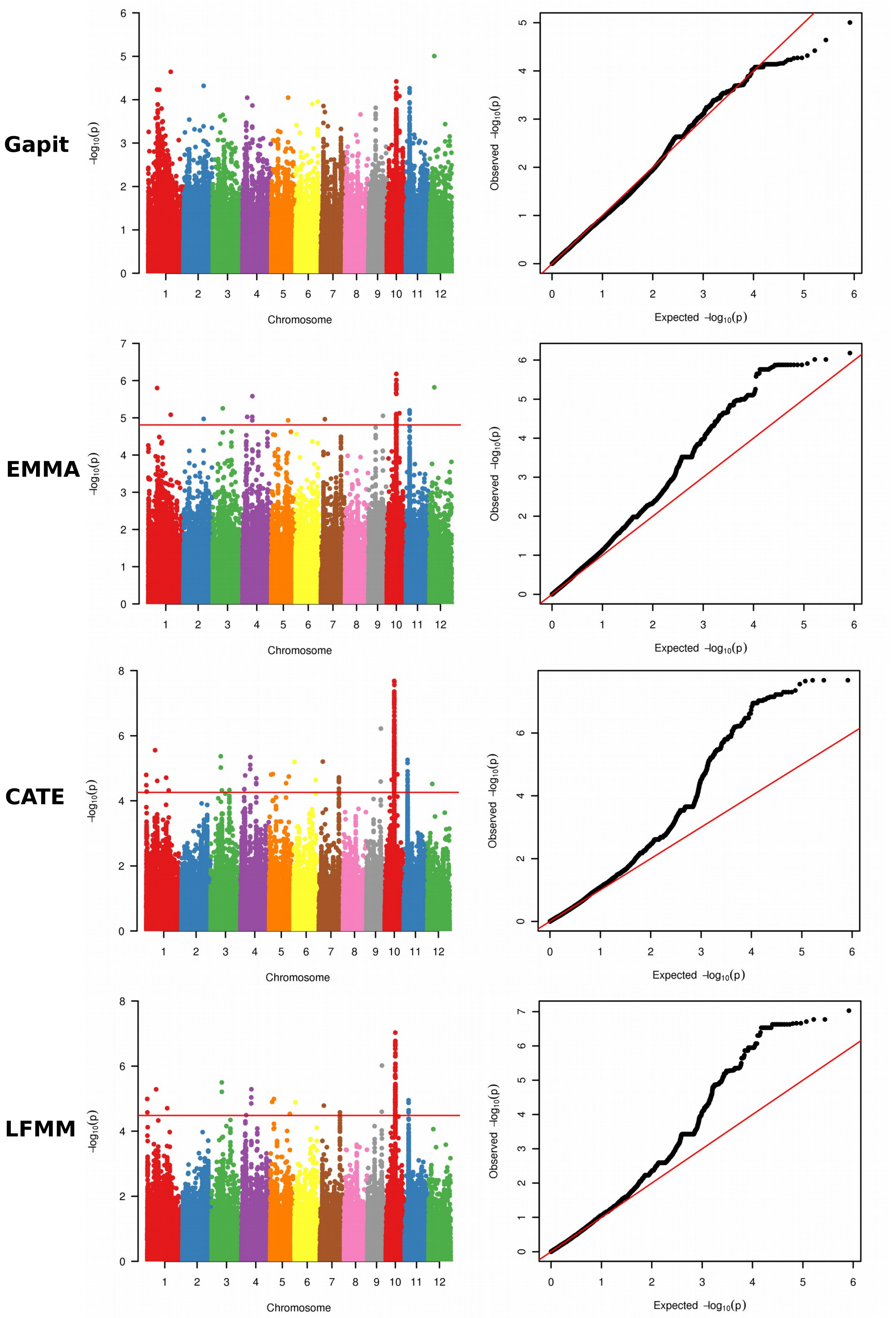

### Additional File 4

#### RYMV resistance

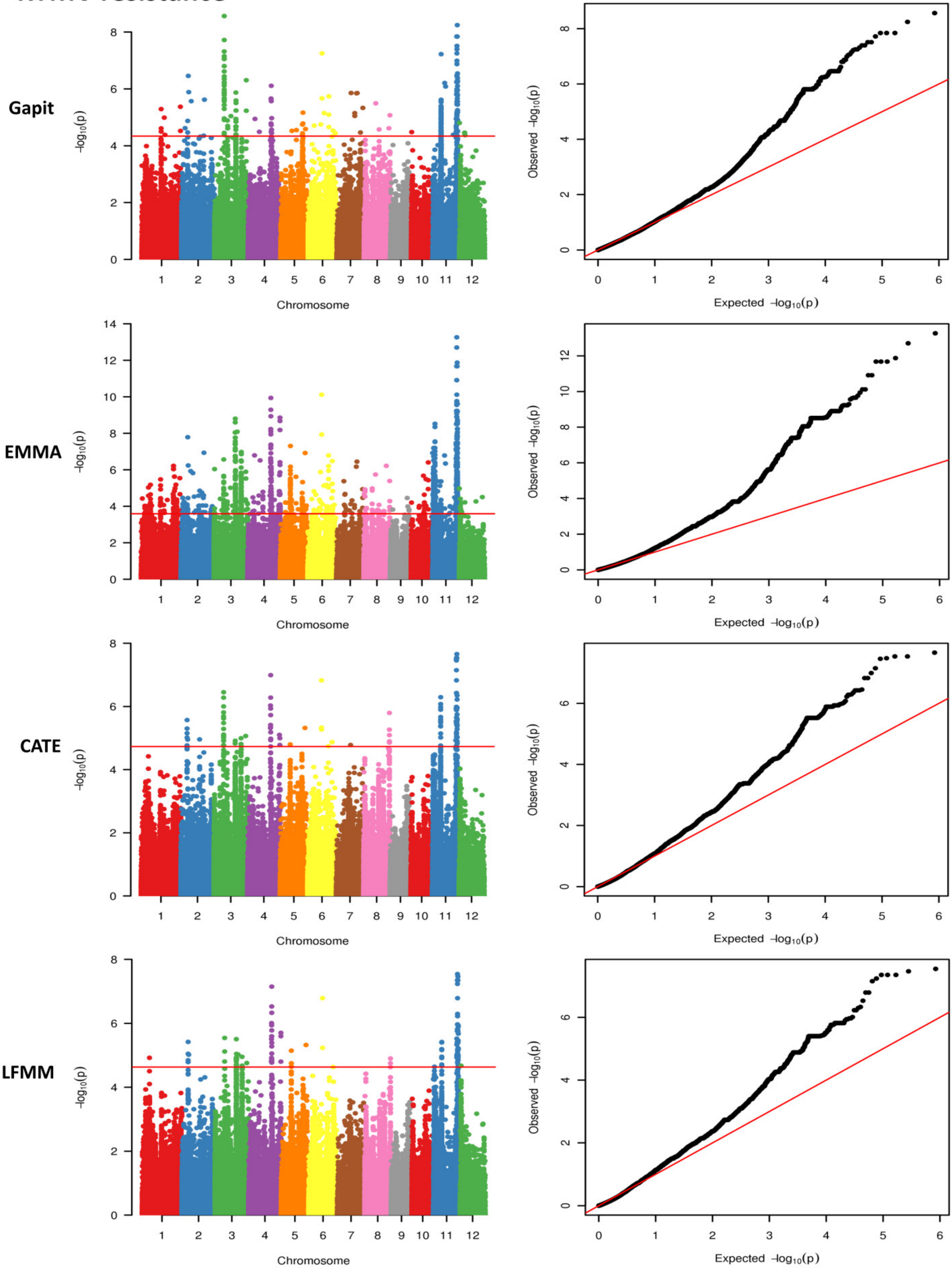
