## Additional File 5 for "Genome wide association study pinpoints key agronomic QTLs in African rice *Oryza glaberrima*"

### BioPC1

**Gapit**

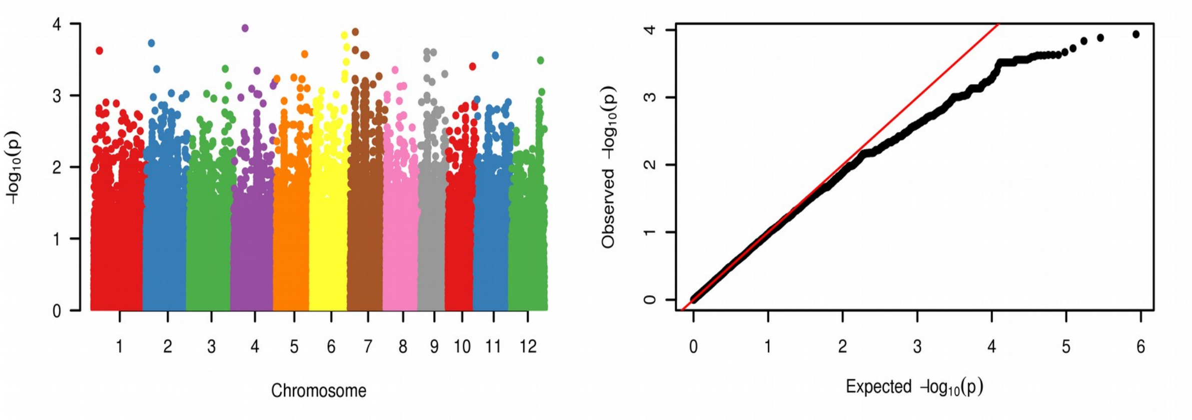


**EMMA**

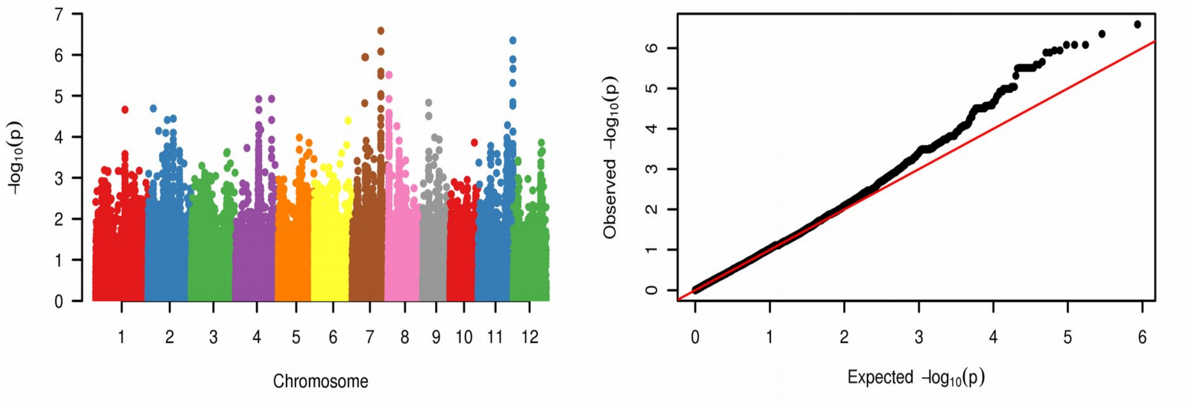


**CATE**

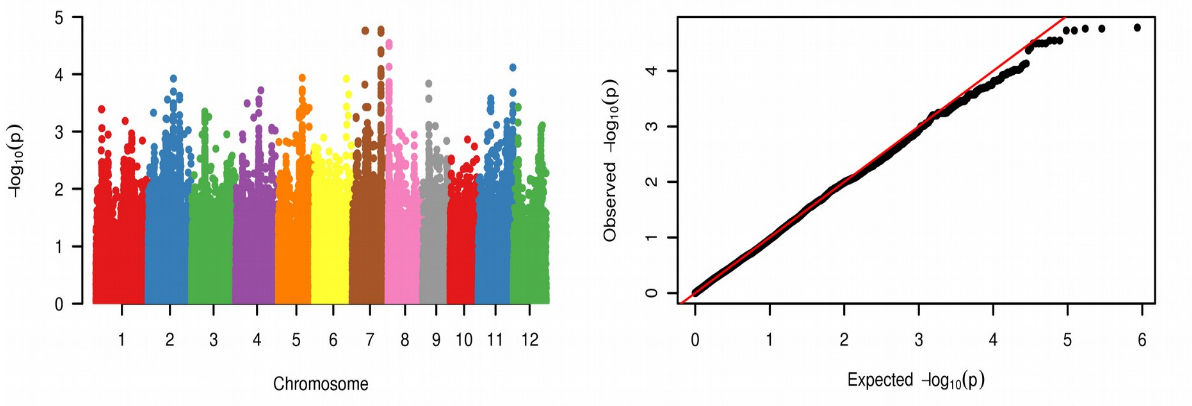


**LFMM**

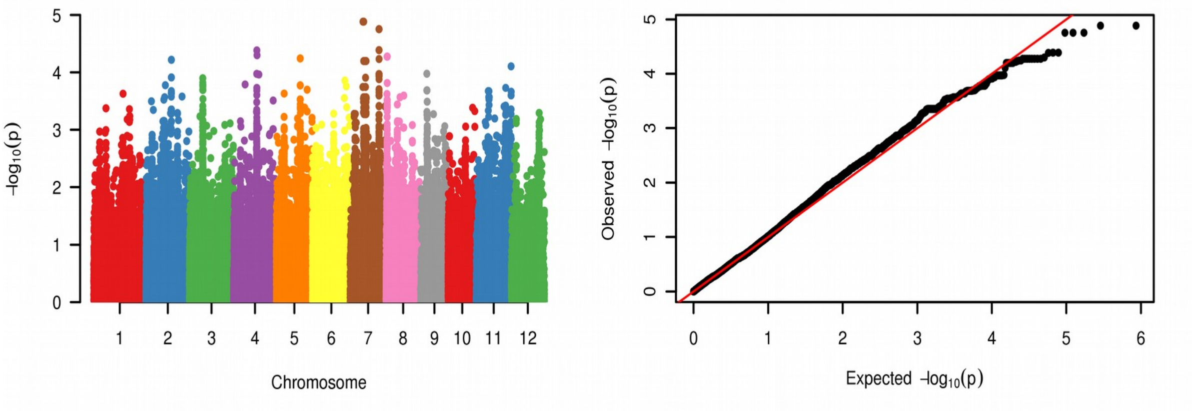

### BioPC2

**Gapit**

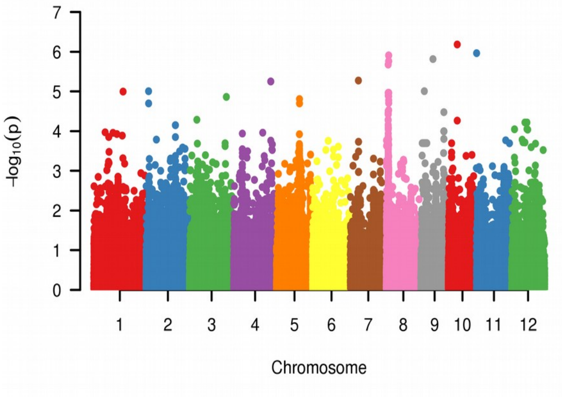


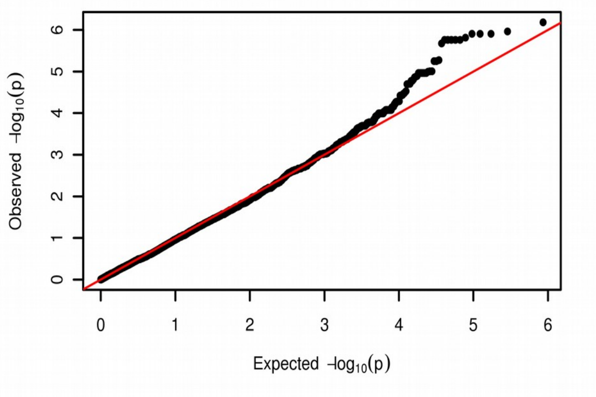


**EMMA**

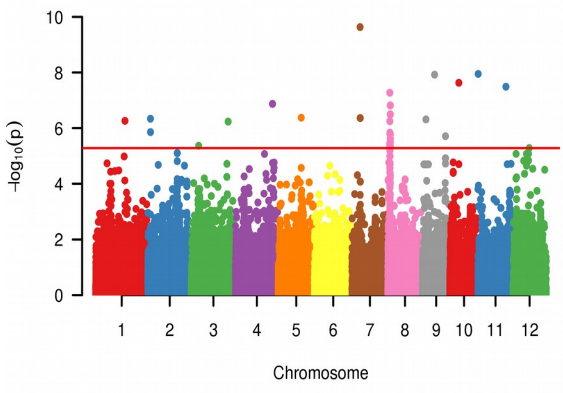


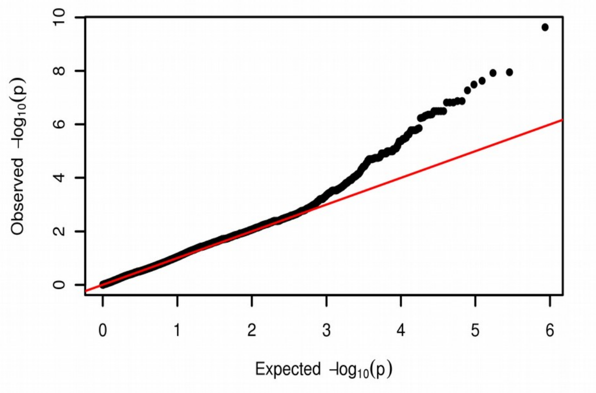


**CATE**

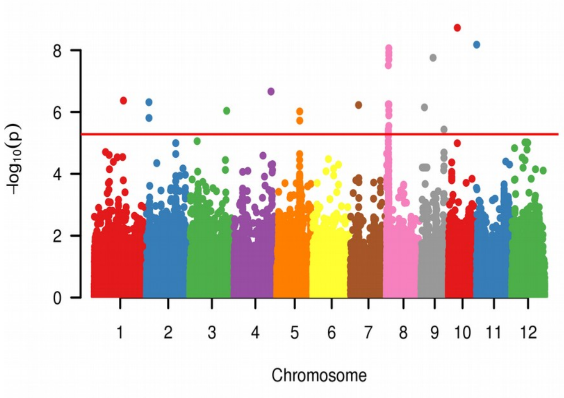


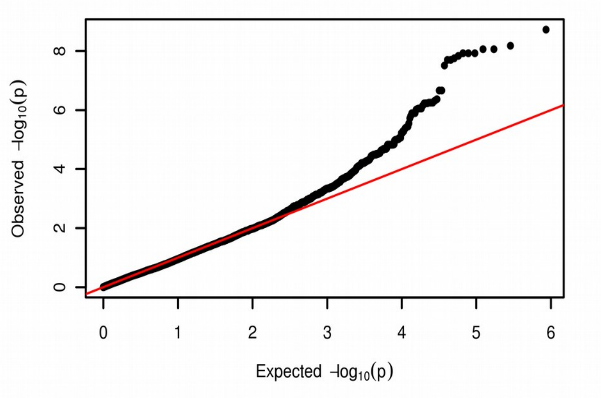


**LFMM**

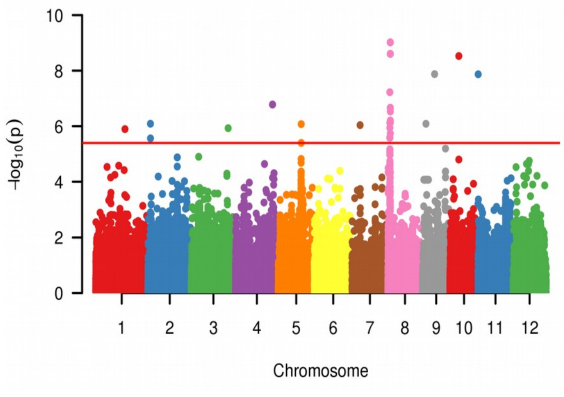


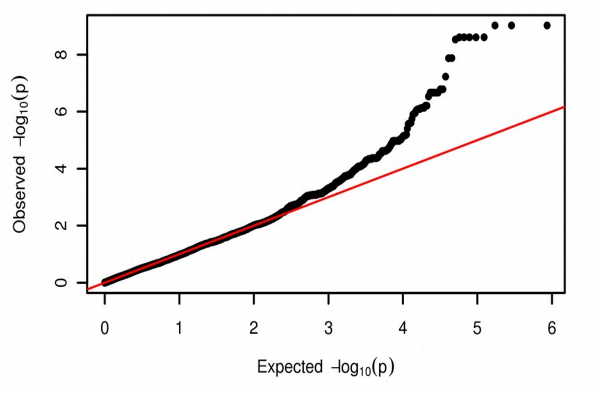

### TmaxPC1

**Gapit**

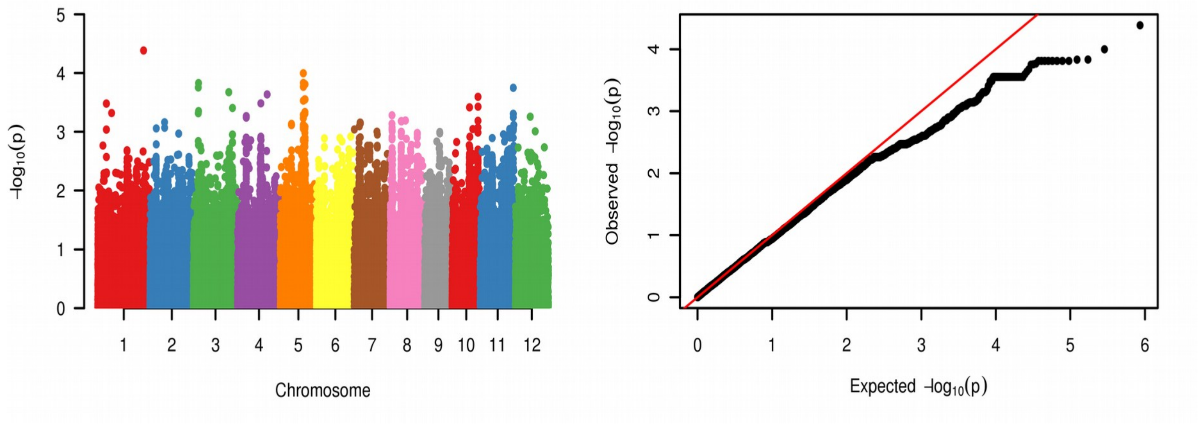


**EMMA**

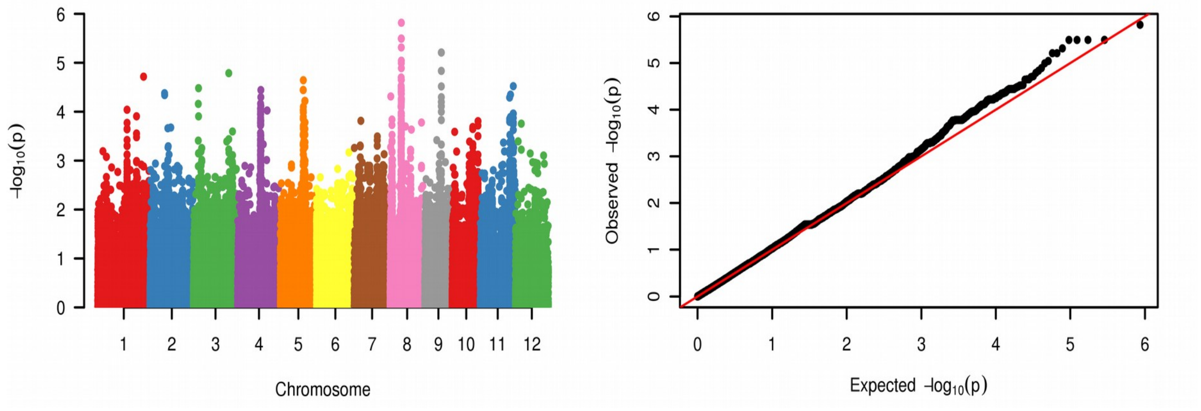


**CATE**

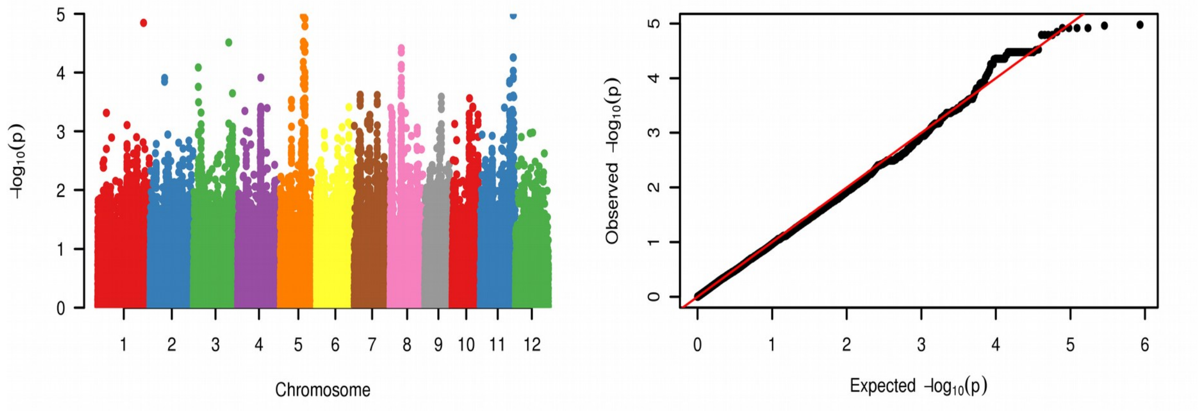


**LFMM**

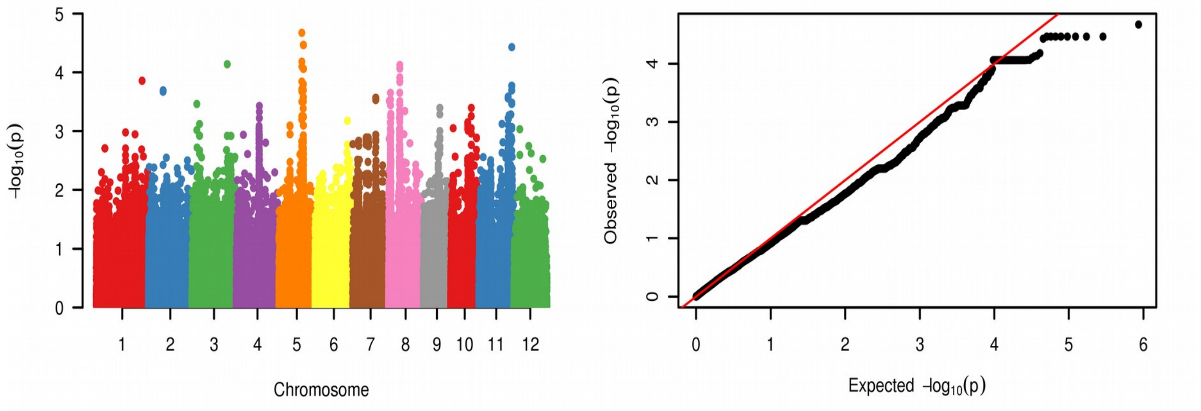

### TmaxPC2

**Gapit**

**EMMA**

**CATE**

**LFMM**

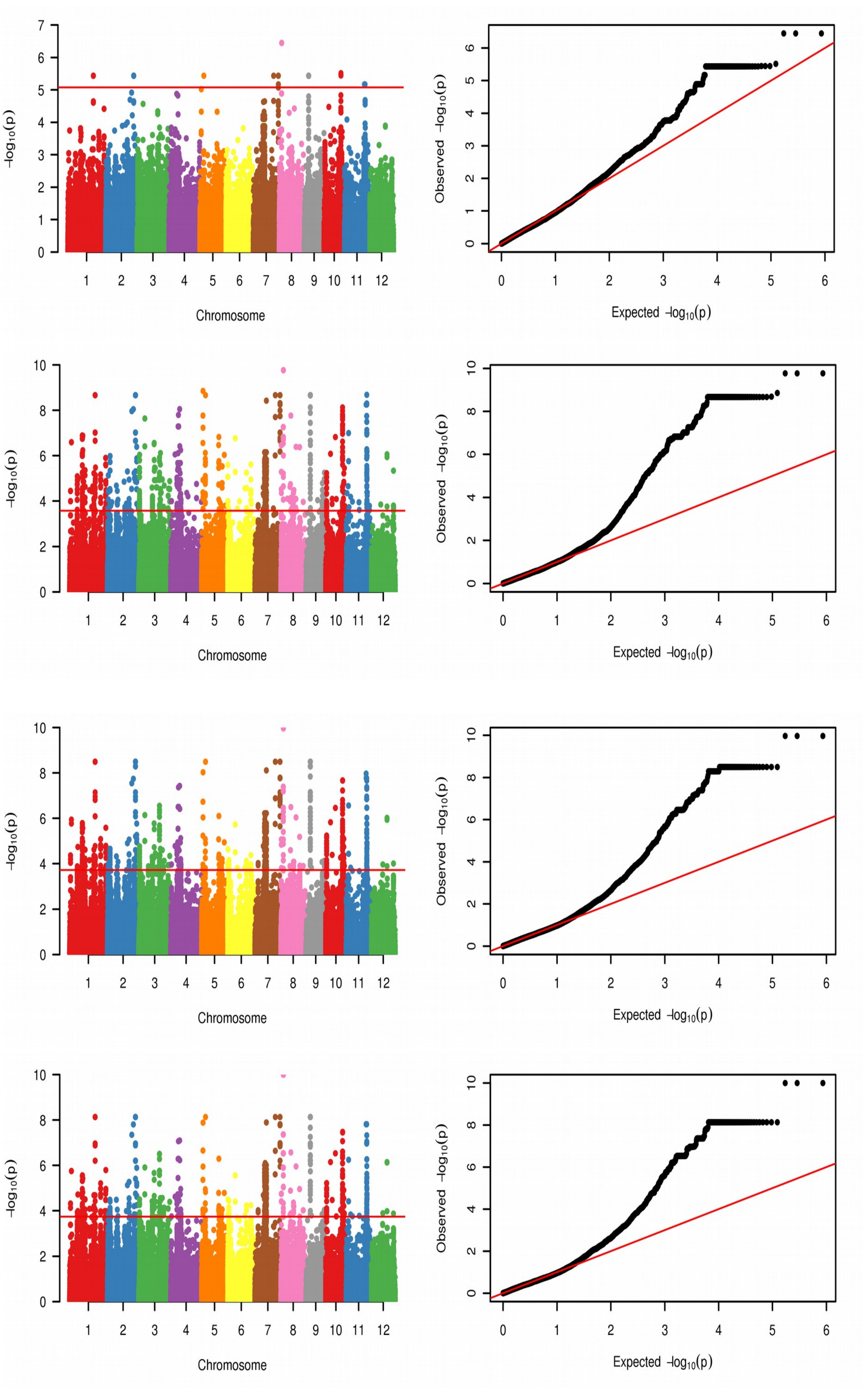
